# Two opposing gene expression patterns within *ATRX* aberrant neuroblastoma

**DOI:** 10.1101/2022.10.25.513663

**Authors:** Michael R. van Gerven, Linda Schild, Jennemiek van Arkel, Bianca Koopmans, Luuk A. Broeils, Loes A. M. Meijs, Romy van Oosterhout, Max M. van Noesel, Jan Koster, Sander R. van Hooff, Jan J. Molenaar, Marlinde van den Boogaard

## Abstract

Neuroblastoma is the most common extracranial solid tumor in children. A subgroup of high-risk patients is characterized by aberrations in the chromatin remodeller ATRX that is encoded by 35 exons. In contrast to other pediatric cancer where *ATRX* point mutations are most frequent, multi-exon deletions (MEDs) are the most frequent type of *ATRX* aberrations in neuroblastoma. Of these MEDs 75% are predicted to produce in-frame fusion proteins, suggesting a potential gain-of-function effect compared to nonsense mutations. For neuroblastoma there are only a few patient-derived *ATRX* aberrant models. Therefore, we created isogenic *ATRX* aberrant models using CRISPR-Cas9 in several neuroblastoma cell lines and one tumoroid and performed total RNA-sequencing on these and on the patient-derived model. Gene set enrichment analysis (GSEA) showed decreased expression of genes related to both ribosome biogenesis and several metabolic process in our isogenic *ATRX* exon 2-10 MED model systems, the patient-derived MED models and in tumor data containing two patients with an *ATRX* exon 2-10 MED. Interestingly, for our isogenic *ATRX* knock-out and exon 2-13 MED models GSEA revealed an opposite expression pattern characterized by increased expression of genes related to ribosome biogenesis and several metabolic process. Our validations confirmed a potential role of ATRX in the regulation of ribosome homeostasis. In this manner we identified two distinct molecular expression patterns within *ATRX* aberrant neuroblastomas with important implications for the need of distinct treatment regimens.

## Introduction

Neuroblastoma is the most common extracranial solid tumor in pediatric cancer and arises during the development of fetal adrenal neuroblasts^1,2^. In recent years, the survival rates have improved to 81% for neuroblastoma in general^3^. However, the survival for high-risk patients remains only 50% despite intensive treatment. Recent data has shown that high-risk patients can be stratified in four genetic subgroups: *MYCN* amplified, *TERT* rearranged, *ATRX* aberrant or none of these three aberrations, in which *MYCN* amplifications and *ATRX* aberrations are mutually exclusive^4^. *ATRX* is a commonly mutated gene in pediatric cancer and its precise molecular role in neuroblastoma development is still unclear.

The chromatin remodeler ATRX is encoded by 35 exons localized on the X-chromosome and is involved in a plethora of nuclear processes. Its most prominent role is the chromatin incorporation of the histone variant H3.3 together with its binding partner DAXX to maintain genomic integrity and a heterochromatin state at pericentromeric and telomeric regions^5–7^. *ATRX* aberrations are associated with Alternative Lengthening of Telomeres (ALT)^8,9^. ALT is a telomerase-independent telomeric maintenance mechanism. This mechanism is a homologous recombination-based process that is still poorly understood. A currently untested theory is that the deposition of H3.3 at telomeric regions is necessary to prevent formation of G-quadruplexes to limit the amount of fork collapse and concomitantly double-stranded DNA breaks (DSBs)^10^. It is suggested that the process of ALT occurs due to the faulty and altered repair of these induced DSBs. Many tumors displaying ALT have *ATRX* aberrations, however several *ATRX* wild-type tumors also display ALT and how ATRX aberration contribute to ALT is currently unknown^11^. Thus, the precise role of ATRX within the development of ALT has not yet been elucidated.

Previously, we have shown that *ATRX* multi-exon deletions (MEDs) are almost exclusively present in neuroblastoma whereas other pediatric cancers are dominated by point mutations^8^. Of these MEDs 75% are predicted to be in-frame and for the most common MEDs it has been shown that these still result in protein production, suggesting a potential gain-of-function effect compared to wild-type. So far 30 unique *ATRX* MEDs have been reported in neuroblastoma, of which only three constitute the far majority of cases^8^. Many of these 30 unique MEDs lack a large part of the N-terminal region including exons 8-9. Furthermore, several rare MEDs have been identified that lack a smaller part of the gene including exons 11-12, which contains the DAXX-binding domain. In neuroblastoma only very few patient-derived models exists and the deletions that are present only represent a small fraction of the deletions that are present in patients, while for nonsense and missense mutations there are no model systems at all. Nonsense, missense and distinct *ATRX* deletions could be molecularly very different and therefore might need distinct therapies.

In order to study the molecular role of different *ATRX* aberrations in neuroblastoma, we created isogenic ATRX knock-out (KO) and several distinct in-frame MEDs, including some rare deletions, in neuroblastoma cell line and organoid models. Gene expression analysis was conducted for all generated models and for three patient-derived *ATRX* MED models. We were able to identify the presence of two opposing molecular expression profiles for different *ATRX* aberrations.

## Results

### Characterization of neuroblastoma cell lines with ATRX multi-exon deletions

To study the role of ATRX aberrations in neuroblastoma development, we acquired two classical neuroblastoma cell lines, SK-N-MM and CHLA-90, and the tumoroid AMC772T2 that we previously established in our lab^4^. We first validated the genomic aberrations of these three *ATRX* MED models using a PCR-based assay on cDNA, since the exact DNA breakpoint within the introns are unknown. For the male cell line CHLA-90 we identified transcripts confirming a genomic deletion of exon 3-9 (Figure 1A), as previously reported ^12,13^. For the female cell line SK-N-MM we confirmed the nonsense mutation (Figure1A; K1367*) as described by Qadeer et al, and we identified transcripts containing exon 2-9 and exon 2-10 MEDs (Figure 1A). This indicates that on the genomic level there is a deletion of exons 2-9, which is predicted to be out-of-frame. However, by skipping exon 10, in-frame transcripts are generated^14^. For the tumoroid AMC772T2 that is derived from a male patient we found both exon 2-9 and exon 2-10 MED transcripts (Figure 1A), indicating an exon 2-9 MED on the genomic level. Thus, our patient-derived models contain *ATRX* MEDs on the genomic and transcriptomic level.

**Figure 1.**
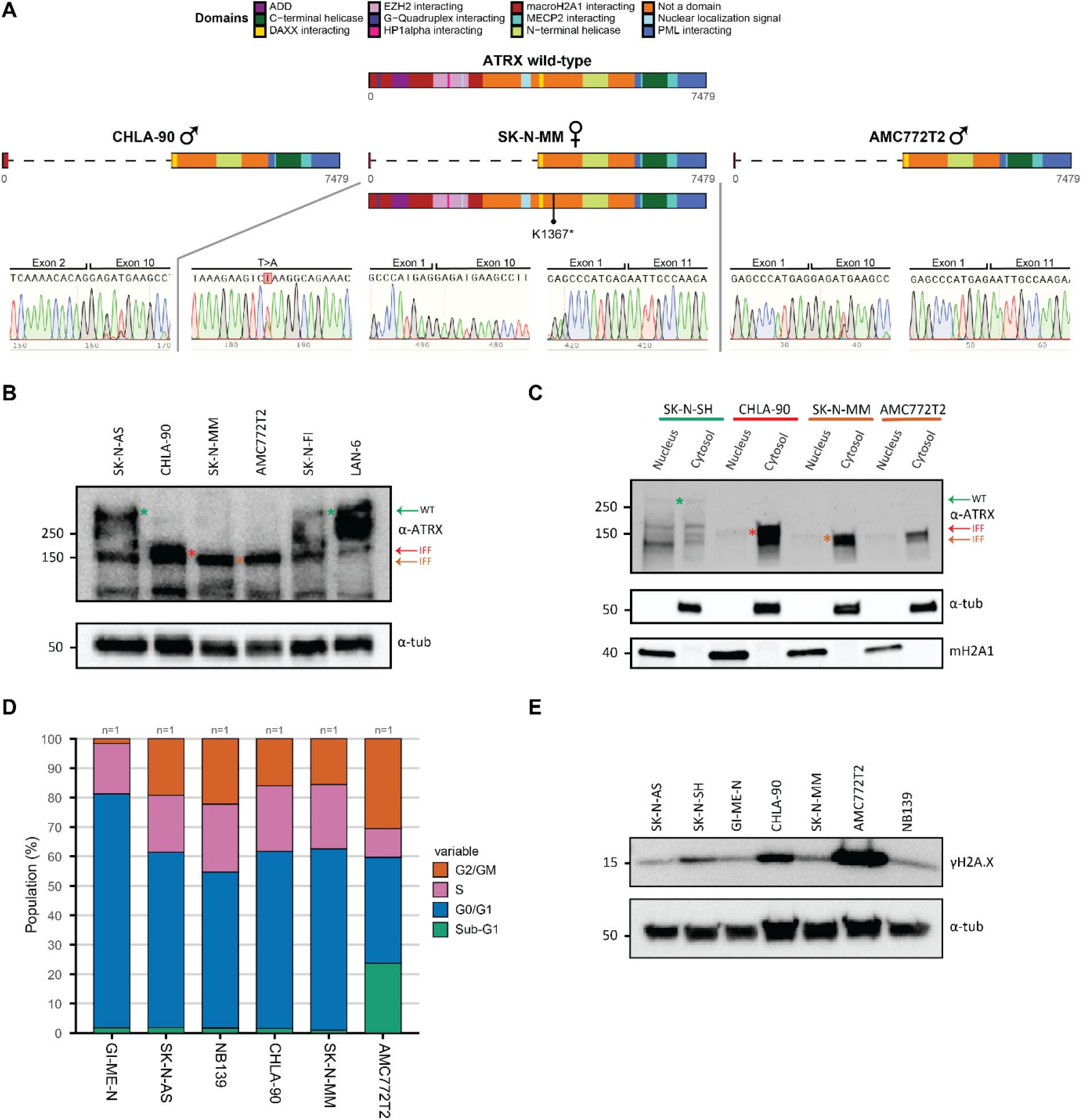
Characterization of neuroblastoma cell lines with ATRX multi-exon deletions. **A)** Genetic confirmation of the *ATRX* aberrations of three patient-derived *ATRX* MED neuroblastoma models (CHLA-90, SK-N-MM and AMC772T2) utilising *ATRX*-targeted cDNA PCR amplification and sequencing. Only for the SK-N-MM point mutation performed validation in gDNA. **B)** Western blot confirming the presence of mutant in-frame fusion (IFF) ATRX protein product in the three patient-derived *ATRX* MED models. **C)** Western blot fractionation experiment showing cytosolic retention of the mutant IFF products. **D)** Cell cycle distribution analysis of *ATRX* wild-type and *ATRX* MED models. “n=” indicates the number of biological replicates used in these experiments. **E)** Western blot of γH2A.X abundance in *ATRX* wild-type and *ATRX* MED models.

Subsequently, we assessed the protein expression of ATRX utilizing an antibody against the C-terminus that recognizes both full-length and in-frame fusion (IFF) ATRX protein products. We detected full-length ATRX protein (280 kDa) for SK-N-AS, LAN-6, and SK-N-FI (control cell lines) and ATRX IFF protein (~140-150 kDa) for CHLA-90, SK-N-MM and AMC772T2 (Figure 1B). In all three *ATRX* models multiple key proteins domains are lost, including the nuclear localization signal (Figure 1A). Therefore, we assessed protein localization using fractionation western blot experiments, in which we detected a strong retention of IFF proteins in the cytosol with minimal amounts in the nucleus (Figure 1C). In contrast, in the ATRX wild-type cell line SK-N-SH we observe equal amounts of full-length protein in both fractions. Lastly, we assessed the localization of ATRX to heterochromatic foci together with its binding partner HP1α that marks pericentric heterochromatic regions^7,15^. In the ATRX wild-type cell line GI-ME-N we observe clear ATRX foci that co-localize with HP1α foci, while in all three patient-derived *ATRX* MED models (PD^ΔATRX^) we observe diffuse ATRX staining and absence of ATRX foci (S1). Thus, ATRX MEDs are expressed on the protein level, but are strongly retained in the cytosol and no longer form ATRX nuclear foci.

ATRX aberrations are strongly associated with ALT^8,16^ and therefore we employed two assays to confirm this ALT phenotype within our PD^ΔATRX^ models. Our ALT-associated PML bodies (APBs) staining confirmed the presence of ALT in all three PD^ΔATRX^ models, in which we observe strong telomeric staining that co-localizes with PML protein (S2). On the telomeric southern blot we observed extremely long and heterogeneous telomeric length in our PD^ΔATRX^ models as well as in two additional ALT models (SK-N-FI and LAN-6, both *ATRX* wild-type), but not in SK-N-SH (S3A). In conclusion, we confirmed the presence of ALT in our PD^ΔATRX^ models.

ATRX is involved in prometaphase to metaphase transition^17^ and in the removal of G-quadruplexes and R-loops^18,19^. These secondary DNA structures hinder progression of replication, which might lead to replication-fork stalling and ultimately in fork collapse and increased DNA damage. Knock-out of *ATRX* has been shown to result in both prolonged mitosis^17^ and S-phase^20^, the latter as a result of increased replication stress. However, we do not observe an increased proportion of cells in S or G2/M phase in our PD^ΔATRX^ models compared to wild-type models (Figure 1D). We also do not observe changes in the rate of proliferation compared to wild-type cells (S3B). Previously, it has been reported that *ATRX* knock-out leads to increased DSBs^21^. However, we only found increased DNA damage in two of the three PD^ΔATRX^ models (Figure 1E). Thus, we were unable to detect ATRX-specific disturbances in cell cycle progression and no clear association between *ATRX* aberrations and the level of DNA damage.

### Generation and validation of ATRX aberrant isogenic cell lines/tumoroids

Currently, only very few *ATRX* aberrant models are available, and they only represent a small fraction of all observed *ATRX* patient aberrations. Therefore, we made isogenic model systems, which have the additional advantage of the presence of a reference mother-line. To create *ATRX* KO models, that represent *ATRX* nonsense mutations, we used a CRISPR guide targeting exon 4 and a plasmid containing homology-arms to knock-in a GFP puromycin construct for selection (Figure 2A). After selection, cells were sorted to generate single-cell clones. To generate large MEDs we used two CRISPR guides targeting the flanking intronic regions (Figure2B) and generated single-cell clones by sorting on GFP-positive cells. We generated the most common MED of exon 2-10, the rare MED of exon 2-13 and the rare MED of exon 10-12 (for more information about all observed MED in literature see ^8^). We attempted to create both *ATRX* KO and *ATRX* MEDs in several cell lines, an overview of our (un)successful attempts can be found in Table S1 and protein confirmation is shown in S4A. Thus, in the end we established a total of 20 *ATRX* aberrant isogenic clones.

**Figure 2.**
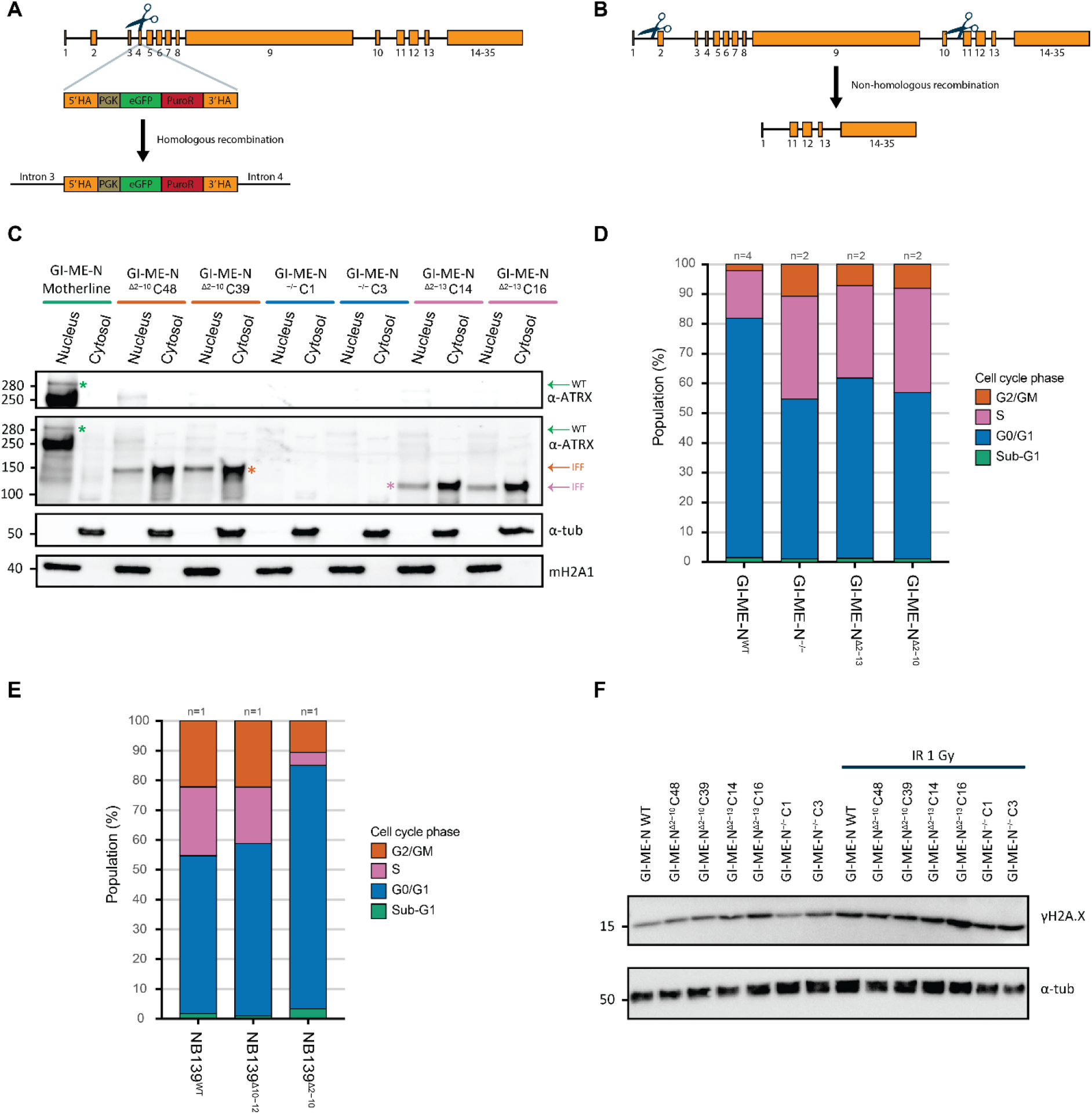
Generation and validation of ATRX aberrant isogenic cell lines/tumoroids. **A)** Overview of the strategy utilised to generate ATRX KO models with CRISPR-Cas9 targeting exon 4. HA: Homology Arms, PGK: promotor, eGFP: Green fluorescent Protein, PuroR: puromycin resistance gene and scissors: guide+Cas9. **B)** Overview of the strategy applied to create ATRX MED models with a dual-guide CRISPR-Cas9 strategy. **C)** Western blot fractionation experiment validating cytosolic retention of the mutant *ATRX* IFF products generated by multiple isogenic GI-ME-N clones. **D)** Cell cycle distribution analysis of *ATRX* isogenic GI-ME-N models. **E)** Cell cycle distribution analysis isogenic NB139 models. **D-E)**“n=” indicates the number of biological replicates used in these experiments. **F)** Western blot of γH2A.X abundance in isogenic *ATRX* aberrant GI-ME-N models at baseline and upon 1 gray radiation (IR 1 Gy).

To further validate the isogenic models, we performed the same experiments as on our PD^ΔATRX^ models. For all our isogenic MED models of exons 2-10 and 2-13 we detected cytosolic retention of ATRX IFF proteins and diffuse ATRX staining (Figure 2C and S6-8), similarly to what we observed in the PD^ΔATRX^ models. Also diffuse ATRX staining was observed for the NB139 exon 10-12 MED model. This might indicate that the loss of the DAXX binding domain in this model in sufficient to disturb ATRX localization to pericentric heterochromatin. In contrast, we could not identify any signs of ALT in our isogenic models by APBs stainings (S9-11) and telomeric southern blot analysis (S4B). This could indicate that ALT is depending on a more complex genomic background and is in line with earlier reports showing that inducing *ATRX* aberrations does not necessarily result in ALT activation ^20,22–24^. In contrast to our PD^ΔATRX^ models, we do observe a larger fraction of cells in both S and G2/M phase and fewer cells in GO/G1 in all our *ATRX* aberrant GI-ME-N models compared to wild-type (Figure 2D). Additionally, we observed unchanged proliferation rates for all *ATRX* aberrant GI-ME-N models (S5B). Suggesting that only the fractions of cells undergoing proliferation increased, but not the rate of proliferation. For our NB139 *ATRX* MED of exon 2-10 we observed changes in both cell cycle and proliferation. However, instead of increased S and G2/M and decreased G0/G1 that we observed for our isogenic *ATRX* aberrant GI-ME-N models, we observed the opposite in our NB139 exon 2-10 MED (Figure 2E). Furthermore, we observe more cells with a stronger violet trace signal compared to wild-type, indicating a decreased rate of cell proliferation (S5C). On the contrary we do not observe any changes in our isogenic SK-N-AS *ATRX* KO models for cell cycle or proliferation (S5A and D). Lastly, we assessed the amount of DSBs in our isogenic models but did not observe increased DNA damage (Figure 2F and S4C), not even upon induction by radiation in our GI-ME-N models (Figure 2F). In summary, our created isogenic models recapitulated some of the phenotypes observed in the PD^ΔATRX^ models, however the effect of *ATRX* aberrations on cell cycle might be cell line/type dependent.

### Strong overlap of differentially expressed genes between ATRX^Δ2-13^ and ATRX^−/−^ GI-ME-N models

To study the molecular landscape of ATRX aberrant neuroblastoma we assessed the transcriptomes by performing total RNA-sequencing. When we performed PCA of all isogenic models we observe separation based on the mother-lines (S12A). From the PCA of GI-ME-N, SK-N-AS and NB139 we can observe nice separation of wild-type versus *ATRX* mutant clones (S12B-D). Only for GI-ME-N^Δ2-13^ clone 15 we observed clustering with the GI-ME-N^−/−^ clones that we could not explain.

We performed differential expression analyses for SK-N-AS^−/−^ (*ATRX* KO), GI-ME-N^−/−^, GI-ME-N^Δ2-10^ (*ATRX* MED exon 2-10), GI-ME-N^Δ2-13^ and NB139^Δ2-10^ compared to corresponding wild-type clones. Several thousands of genes were differentially expressed in each of the 5 expression analyses (Table S2). To determine if the different *ATRX* aberrations resulted in similar changes in gene expression we took the overlap of both the down and upregulated genes within the GI-ME-N models. We noticed a striking overlap between GI-ME-N^−/−^ and GI-ME-N^Δ2-13^ and very little overlap with GI-ME-N^Δ2-10^, indicating that there might be two distinct expression patterns within *ATRX* aberrant GI-ME-N models (Figure 3A). Therefore, suggesting one expression pattern related to complete inactivation of ATRX (GI-ME-N^−/−^ and GI-ME-N^Δ2-13^) and one related to the IFF with remaining or gained protein activity (GI-ME-N^Δ2-10^).

**Figure 3.**
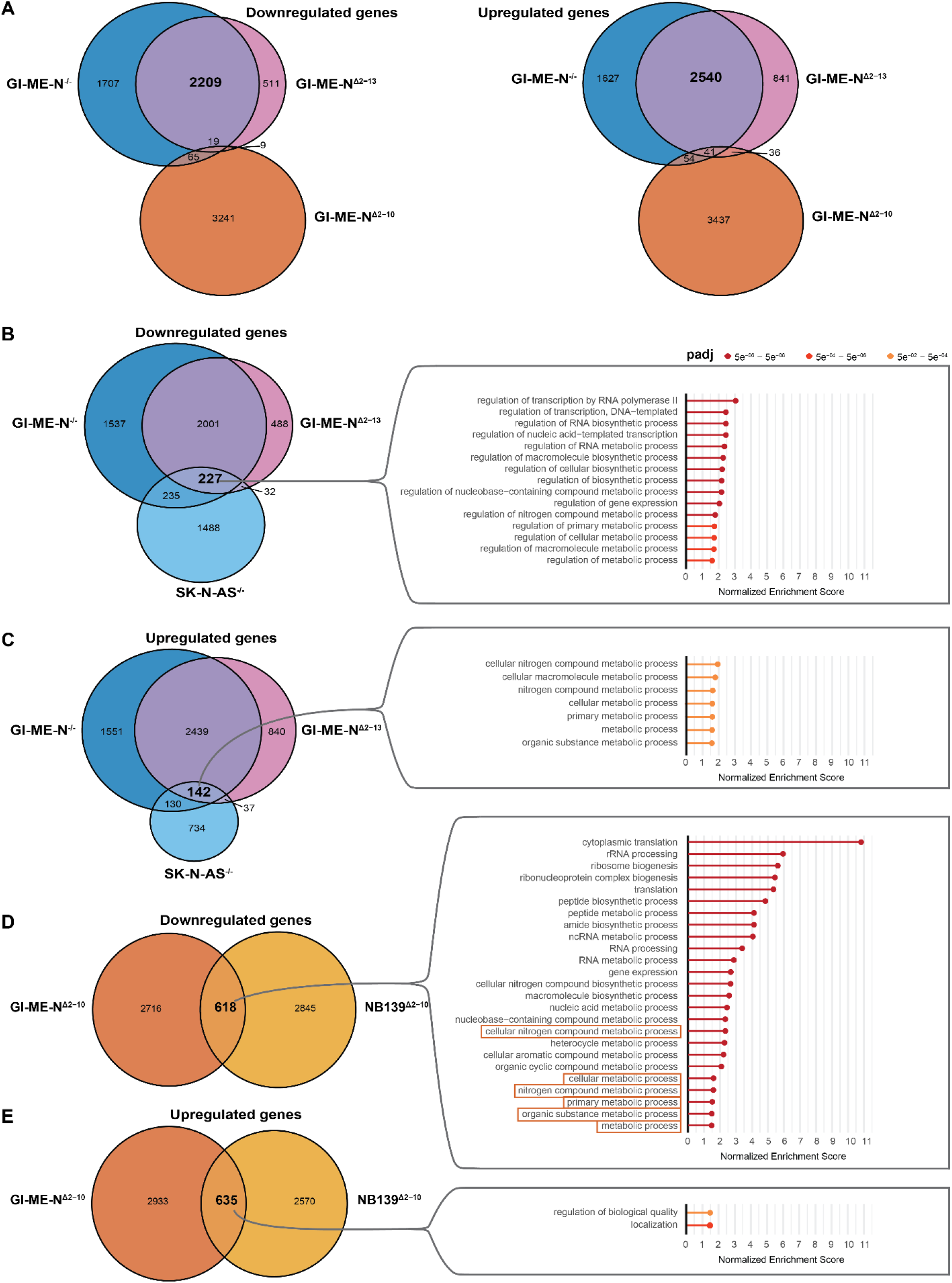
Overlap of differentially expressed genes (DEGs) and Gene Ontology (GO) analysis identifies two distinct gene expression patterns within *ATRX* aberrant neuroblastoma. **A)** Overlapping DEGs for both the down and upregulated genes within the isogenic *ATRX* aberrant GI-ME-N models. **B)** GO analysis of the overlapping downregulated DEGs between all *ATRX^−/−^* and *ATRX*^Δ2-13^ isogenic models. **C)** GO analysis of the overlapping upregulated DEGs between all *ATRX^−/−^* and *ATRX*^Δ2-13^ isogenic models. **D)** GO analysis of the overlapping downregulated DEGs between both *ATRX*^Δ2-10^ isogenic models. Only the top 25 significant gene ontologies are shown here, for all significant GO see S13. Orange boxes highlighted the same terms as observed in figure 3C. **E)** GO analysis of the overlapping upregulated DEGs between both *ATRX*^Δ2-10^ isogenic models.

### Gene ontology reveals increased expression of genes related to metabolic process in ATRX^Δ2-13^ and ATRX^−/−^ models and decreased expression in ATRX^Δ2-10^ models

To further study this hypothesis, we looked at the overlapping down- and upregulated genes between all KO and *ATRX*^Δ2-13^ models and performed gene ontology (GO) analysis on those genes. GO analysis of the 227 overlapping downregulated genes showed enrichment for the regulation of several RNA and metabolic processes (Figure 3B), while for the upregulated genes we found an overlap of 142 genes that are enriched for seven processes involved in metabolism (Figure 3C). In contrast, GO analysis of the 618 overlapping downregulated genes for our *ATRX*^Δ2-10^ models identified these same seven metabolic processes and also several other terms involved in other metabolic and RNA processes (Figure 3D and S13). While GO analysis of the 635 overlapping upregulated genes for our *ATRX*^Δ2-10^ models showed enrichment for process involved in biological quality and localization (Figure 3E). In all four comparisons between the overlapping differentially expressed genes (Figure 3B-E), we also noticed many non-overlapping genes that are only differentially expressed within one cell line model. This might be explained by the fact that ATRX is a chromatin remodeller and that the effects of *ATRX* aberrations are highly dependent on the epigenetic landscape that is already present in the distinct cell lines.

ATRX is known to bind at the 3’ exon of zinc finger genes ^25^ were it could potentially modulate their expression. Therefore, we performed panther protein class analysis for the overlapping up and down-regulated genes for both the *ATRX* ^−/−^ and *ATRX*^Δ2-13^ models and for the *ATRX*^Δ2-10^ models. Only for the downregulated genes of both we found significant terms, namely zinc finger transcription factors for our *ATRX* ^−/−^ and *ATRX*^Δ2-13^ models (S12G) and proteins involved in RNA processes and translation in *ATRX*^Δ2-10^ models (S12H). Altogether, we can conclude that two distinct expression profiles are present within *ATRX* aberrant neuroblastoma that seem to lead to opposing changes in metabolic processes.

### GSEA reveals two opposing expression patterns within *ATRX* aberrant neuroblastoma

To acquire more understanding of the changed biological processes we performed Gene Set Enrichment Analysis (GSEA) for all isogenic model systems. Additionally, we also performed GSEA for the comparison of PD^ΔATRX^ models with five non-*MYCN* amplified neuroblastoma cell lines (S12E, for number of DEGs see Table S2) and for the comparison of neuroblastoma tumors from the individualized THERapy (iTHER) project. The iTHER data contains 2 tumors with an *ATRX* MED of exon 2-10 (iTHER^Δ2-10^) and 7 *ATRX* wild-type and non-*MYCN* amplified tumors (S12F, for number of DEGs see Table S2). Thus, in total we performed 7 GSEA for both the gene set database gene ontology biological process (GO BP) and Reactome (Figure 4A). The majority of neuroblastoma patients have a MED of exon 2-10^8^ and therefore we visualized all the overlapping significantly changed genes sets with the same Normalised Enrichment Score (NES) directionality (e.g. – or +) between GI-ME-N^Δ2-10^, NB139^Δ2-10^, PD^ΔATRX^ and iTHER^Δ2-10^ in bubble plots (Figure 4A). Our bubble plot of all the overlapping significant GO BP gene sets shows decreased expression of genes related to ribosome biogenesis, translation, and metabolism in these models (Figure 4B, S14B, E-G). While we observe the complete opposite patterns for GI-ME-N^−/−^, SK-N-AS^−/−^ and GI-ME-N^Δ2-13^, which is in line with our GO analysis results (Figure 4B, S14A, C-D). This pattern is confirmed by the comparisons of the GSEA of Reactome (Figure 4C). Taken together, this suggests two opposing expression profiles within *ATRX* aberrant neuroblastoma.

**Figure 4.**
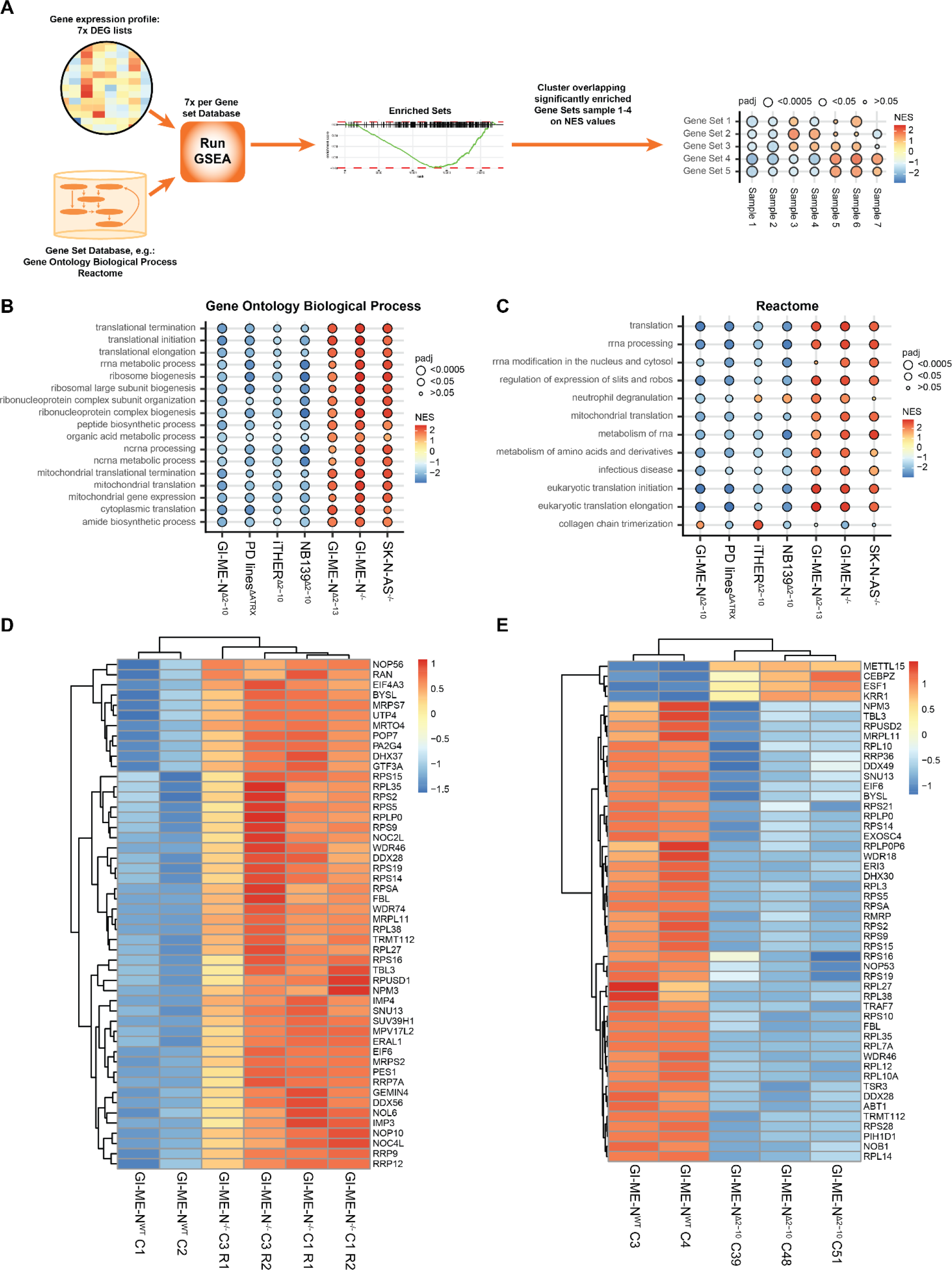
Gene Set Enrichment Analysis (GSEA) reveals two opposing expression patterns within *ATRX* aberrant neuroblastoma related to ribosome biogenesis. **A)** Overview of the strategy to combine the 7 GSEA for our 7 differentially expressed gene (DEG) lists. NES: Normalised Enrichment Scores. **B)** Heatmap of the correlations between the different GSEA for the gene ontology biological process (GO BP) database, excluding those gene sets that were non-significant in all 7 GSEA. **C-D)** Bubble plot showing all gene sets that were both significant and showed the same directionality of the NES values (positive or negative) in GI-ME-N^Δ2-10^, NB139^Δ2-10^, PD^ΔATRX^ and iTHER^Δ2-10^ for **C)** the GO BP gene set database or **D)** the Reactome gene set database. **E-F)** Heatmap of the top 50 differentially expressed ribosome biogenesis genes for **E)** the isogenic GI-ME-N^−/−^ clones and for **F)** the isogenic GI-ME-N^Δ2-10^ clones. Heatmaps show expression values that were normalized across all samples by Z-score. Both row and column clustering were applied, and distinct clusters were identified.

In the paper of McDowell et al., 1999 it was reported that ATRX binds to the short arm of acrocentric chromosomes were rDNA copies are localised^26^,which together with our GSEA data might suggest a role of ATRX in ribosomal biogenesis. Interestingly, we also observed multiple GO BP gene sets in our GSEA related to cytoplasmic translation, mitochondrial translation, and metabolism (Figure 4B). These GO BP gene sets are all dependent on the abundance of ribosomes, since a reduction or increase in the amount of ribosomes leads to reduced or increased translation capability^27^ and consecutively might lead to changed metabolism. Visualization of the top 50 differentially expressed ribosome biogenesis genes for all our isogenic models and cell line data shows changed expression of both small and large ribosomal proteins and also of many other proteins involved in ribosome biogenesis (Figure 4D and E, S15A-D). Altogether, this suggests that IFFs in *ATRX* may lead to modulations of ribosome homeostasis.

### ATRX is involved in ribosome biogenesis by modulating rRNA expression

MYCN, c-MYC and the ATRX binding partner EZH2 are all directly involved in regulating ribosome biogenesis^28–30^. To exclude an indirect effect of ATRX on ribosome biogenesis via the expression of these genes we assessed their protein abundance. For both MYCN and c-MYC we observe unchanged expression in the isogenic model systems, while for EZH2 we observe a slight decreased in expression in the GI-ME-N^Δ2-10^ clones and a strong decrease in the NB139^Δ2-10^ clone (S16A). However, we also observe decreased EZH2 protein expression in SK-N-AS^−/−^ clone 14. Additionally, we assessed the gene expression of the REST gene, which was previously reported to be overexpressed in CHLA-90 and SK-N-MM compared to LAN-6 and SK-N-FI^31^. However, we did not observe this in both our cell line and tumor data (S16B-C). Thus, altogether we can conclude that these three major genes regulating ribosome biogenesis can be excluded.

As previously mentioned, ATRX binds to the short arms of acrocentric chromosomes were rDNA copies are localized^26^ and therefore it could be involved in modulating the chromatin landscape at these regions and in that manner be involved in rRNA expression regulation. We assessed the rRNA expression in our isogenic models and in the PD^ΔATRX^ models by performing qPCRs on the unspliced pre-rRNA 47S. We observe increased rRNA expression for GI-ME-N^−/−^, SK-N-AS^−/−^ and GI-ME-N^Δ2-13^ (two sample t-test assuming unequal variance: p = 0.015, p = 0.0245, p = 0.177, respectively) and decreased rRNA expression in GI-ME-N^Δ2-10^, NB139^Δ2-10^ and PD^ΔATRX^ ((two sample t-test assuming unequal variance: p = 0.354, p = 0.029, p = 0.0011, respectively) models (Figure 5A-D). This data confirmed a potential role of *ATRX* in ribosome biogenesis, since the changes in rRNA expression we observe is in line with the pattern observed in our GSEA. Lastly, we also wanted to assess the rRNA expression in two CHLA-90 clones with doxycycline inducible *ATRX* wild-type expression. These clones still expressed the IFF protein, and this expression was unchanged upon doxycycline induction (S16D), while *ATRX* wild-type expression was only detectable upon induction (S16E). We observed decreased rRNA expression upon doxycycline induction (Figure 5E; two sample t-test assuming unequal variance, p = 0.068). In conclusion, ATRX is likely involved in ribosome biogenesis and *ATRX* abrogation leads to changed rRNA expression.

**Figure 5.**
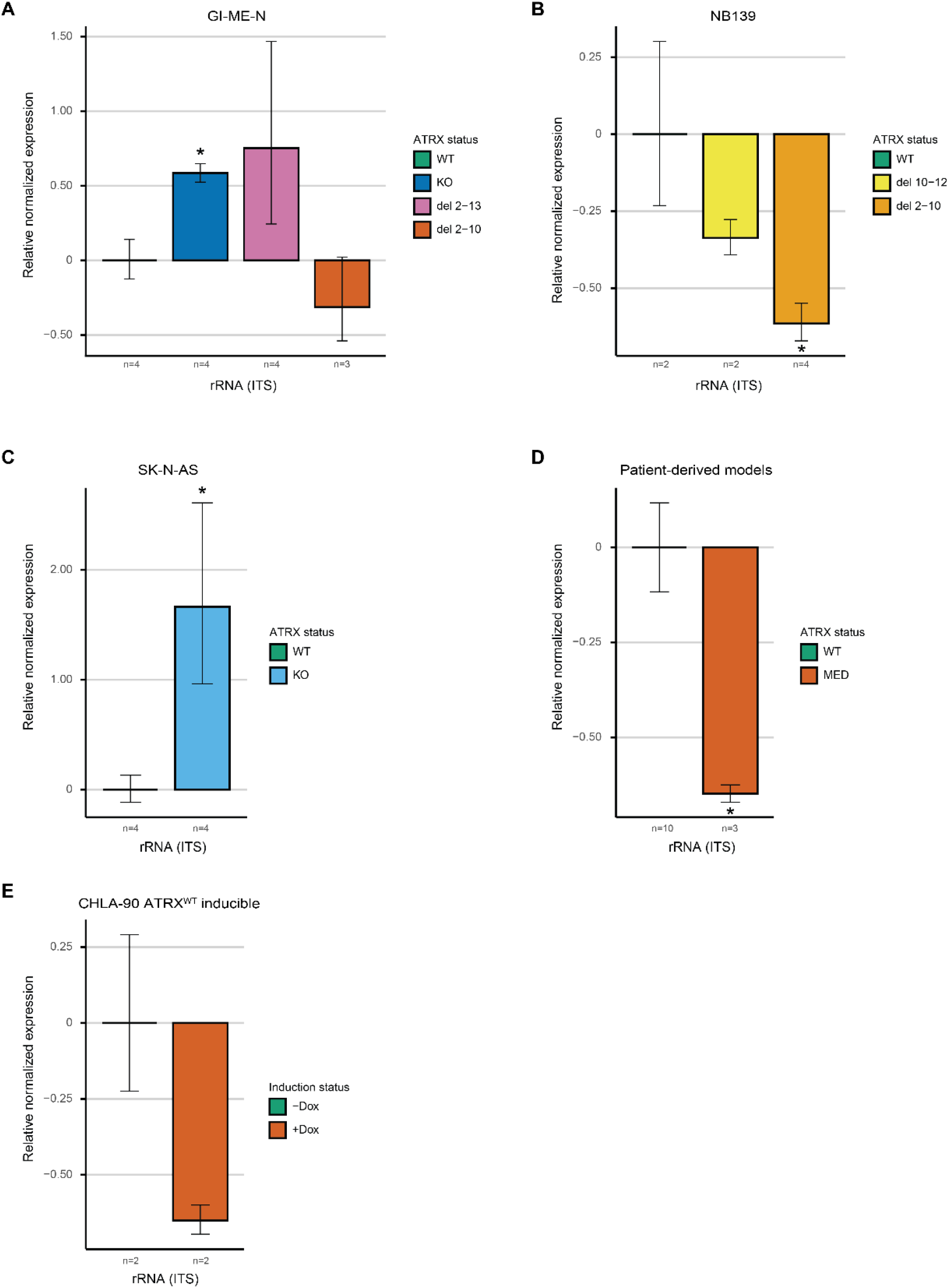
ATRX is involved in ribosome biogenesis by modulating rRNA expression. qPCR-validation on the unspliced 47S pre-RNA using primers for ITS for **A)** the isogenic GI-ME-N clones. **B)** the isogenic NB139 clones. **C)** the isogenic SK-N-AS clones. **D)** the patient-derived *ATRX* MED models. **E)** the isogenic *ATRX*^WT^ doxycycline-inducible CHLA-90 clones. **A-E)**“n=” indicates the number of biological replicates used and for each of these biological replicates three technical replicates were used.

## Discussion

The aim of this study was to assess whether different *ATRX* aberrations are molecularly distinct from one another and how they might contribute to tumor development. We developed a total of 20 isogenic clones of several distinct *ATRX* aberrations. Our RNA analysis revealed a strong overlap in gene expression between *ATRX*^Δ2-13^ and *ATRX*^−/−^ and very little overlap with *ATRX*^Δ2-10^ models. Moreover, we found opposing expression patterns between the *ATRX*^Δ2-13^ and *ATRX*^−/−^ aberrations compared to the *ATRX*^Δ2-10^ aberrations for ribosome biogenesis, ncRNA processes and several metabolic processes. Lastly, we confirmed a potential role of *ATRX* in ribosome biogenesis.

*ATRX* aberrations are strongly associated with ALT and so far all tested neuroblastomas with *ATRX* aberrations utilize this telomere maintenance mechanism^8^. However, we did not observe ALT in any of our *ATRX* aberrant isogenic model systems. The cell lines and tumoroids in which we attempted to make *ATRX* aberrations were all telomerase-dependent and according to literature telomerase-dependent telomeric maintenance could be favored over ALT^32,33^. Therefore, we also attempted to KO *TERT* or *TERC* to force cells in using ALT, but no cells survived and could therefore indicate that other factors might be necessary in conjunction to induce ALT (data not shown). There is indeed evidence that inducing aberrations within the *ATRX* gene does not necessarily cause ALT in different cell types and cancers^20,22–24^.

ATRX has been reported to be involved in cell cycle progression, since *ATRX* KO resulted in prolonged mitosis^17^ and S-phase^20^, the latter as a result of increased replication stress. Interestingly, we observed this for all our *ATRX* aberrant isogenic GI-ME-N models, but not for our isogenic SK-N-AS and NB139 models. Additionally, we also didn’t observe this in the PD^ΔATRX^ models, compared to several wild-type models. The absence of changed cell cycle in our SK-N-AS *ATRX*^−/−^ models could potentially be attributed to the fact that this cell line is *TP53* mutant, which is also to case for two of the three PD^ΔATRX^ models. TP53 is a major regulator of cell cycle progression and is both involved in G1/S and G2/M transition^34^ and therefore *TP53* aberrations can lead to removal of cell cycle blockage. This could therefore mask the effect of the *ATRX* KO on cell cycle in both the SK-N-AS models and the two PD^ΔATRX^ models. An alternative explanation, for the various outcomes of *ATRX* aberrations on cell cycle that we observed, could be that the effect of *ATRX* aberrations is highly cell line specific due to the presence of different chromatin and/or genetic environments.

Changed ribosome biogenesis, proliferation and metabolic process are observed in many cancers^35^. In this study we identified a dichotomy in ribosome biogenesis and several related metabolic processes between the *ATRX*^Δ2-10^ and the *ATRX*^Δ2-13&*−/−*^ (*ATRX*^Δ2-13^ and *ATRX^−/−^)* neuroblastoma tumor cells. Additionally, we show that rRNA expression is downregulated in *ATRX*^Δ2-10^ models and upregulated in *ATRX*^Δ2-13&*−/−*^ models, completely in accordance with our RNA expression data. This suggests that ATRX is in involved in ribosome homeostasis either through direct or indirect modulation of ribosome biogenesis. A hint of the involvement of ATRX in ribosome biogenesis dates from 1999, when it was discovered that ATRX binds to rDNA arrays during metaphase^26^. Only very recently it was observed that ATRX binds to several proteins directly involved in ribosome biogenesis^36^. Another report showed that ATRX binds to the promotor region of rDNA and that in gliomas with nonsense mutations increased ribosome biogenesis can be observed, which is in line with our generated *ATRX* KO models^37^. Intriguingly, for the majority of our created models we did not observe changed proliferative rates, however we observed several altered metabolic processes in our data. This could suggest that the changes in ribosome biogenesis did not reach the threshold to change the proliferation rate but could have reached the threshold to rewire metabolism. There are recent indications that changes in ribosome biogenesis can directly modulate metabolism^38^.

In the majority of cancers high rates of ribosome biogenesis are observed and contribute to tumorigenesis^39^. However, for our *ATRX*^Δ2-10^ models we observe downregulation of ribosome biogenesis suggesting that this will lead to reduced tumorigenesis. Nevertheless, decreased ribosome biogenesis has also be shown to promote tumorigenesis as is observed for patients with ribosomopathies that are prone to the development of certain tumor types^39^. It is postulated that the lower amounts of ribosomes leads to competition between various mRNAs and that tumor suppressor encoding mRNA with lower ribosomal binding affinity could lead to reduced expression of certain tumor suppressors^39^. Also, recent evidence shows that decreases and increases of specific ribosomal proteins are advantageous for tumor development and that these effects are highly cell type and tissue specific^38^. Hopefully, further research will identify and illuminate the precise role(s) of ATRX in ribosome biogenesis.

In conclusion, in this study we successfully established multiple isogenic *ATRX* aberrant models with several distinct *ATRX* aberrations. Utilizing these models, we identified two opposing expression patterns within *ATRX* aberrant neuroblastoma and implicate a potential role of *ATRX* in ribosome biogenesis. Lastly, we want to emphasize that the observed dichotomy in expression pattern within *ATRX* aberrant neuroblastoma suggests the potential need for two distinct treatment regimens, since the *ATRX*^Δ2-10^ and the *ATRX*^Δ2-13&*−/−*^ tumor cells are molecularly very distinct and are therefore likely to respond differently to the same treatment regimen.

## Material and methods

### Cell culture

The neuroblastoma cell lines and tumoroids were cultured in various culture media (Table S3). Originally, we grew NB139 in TIC medium, however at some point our isogenic *ATRX* aberrant NB139 models stopped growing in this medium and therefore we switched to organoid medium with 20% human plasma for both the *ATRX* aberrant and wild-type NB139 clones (wild-type clones had no problem growing in TIC medium; Table S3). All cells were grown in an incubator at 37 °C and 5% CO_2_. Cell lines and tumoroids were routinely checked for mycoplasma infections and authenticated through short tandem repeat profiling.

### sgRNA design and plasmid generation

sgRNAs were designed using the CRISPOR design tool^40^ (sgRNA sequences are listed in Table S4) and were cloned into the pSpCas9(BB)-2A-GFP (PX458) (this plasmid was a gift from Feng Zhang^41^, addgene plasmid #48138). The cloning of the sgRNAs and the two homology arm plasmids (one for ATRX_KO_sgRNA_1 and one for ATRX_KO_sgRNA_2) were performed as described by Boogaard et al^42^. The primers used for amplification of the homology arms and the PGK-eGFP-puromycin cassette are listed in Table S5.

For cloning of the PiggyBac doxycycline-inducible mCMV-Kozak-ATRX-wildtype plasmid we first performed a PCR on cDNA using primers SalI_Kozak_ATRX_FW and ATRX_cDNA_RV (Table S5). Next, we PCR amplified a mCMV from a plasmid using primers XhoI_attL1_mCMV_FW and SalI_mCMV_RV followed by T7 ligation (NEB) of both PCR products. The resulting product was cloned into a pJET1.2/blunt vector (cloneJet PCR cloning Kit, Thermo Scientific, K1231). Subsequently, the resulting plasmid and the IF-GFP-ATRX plasmid (a gift from Michael Dyer^43^, addgene plasmid #45444; exon 6 absent) were digested with XhoI (Promega) and SpeI (Promega) and ligated to generate a Attl1-mCMV-Kozak-ATRX plasmid including exon 6. Next, PCR amplification was performed on the IF-GFP-ATRX plasmid using primers BstEII_ATRX_FW and MluI_Att2L_RV and the resulting product was cloned into a pJET1.2/blunt vector. Subsequently, the resulting plasmid and the Attl1-mCMV-Kozak-ATRX plasmid were digested with BstEII-HF (NEB) and MluI-HF (NEB) and ligated together. The resulting Attl1-mCMV-Kozak-ATRX-Attl2 plasmid was digested using KasI (NEB) and the PB-TAC-ERN plasmid (a gift from Knut Woltjen, addgene plasmid #80475) was digested with MluI followed by a LR gateway clonase reaction (ThermoFisher, 11791020) for 18 hours. All bacterial plates for the cloning of the PiggyBac doxycycline-inducible mCMV-Kozak-ATRX-wildtype plasmid were grown at room.

### Establishing isogenic models (transfections and clone selection)

*ATRX* knock-out clone were established by transfecting SK-N-AS and GI-ME-N with the Cas9 expressing vector containing ATRX_KO_sgRNA_2 (targeting exon 4) and the corresponding homology arm plasmid using Fugene HD transfection reagent (E2312, Promega). Three days after transfection, medium containing 1.5 ug and 1 ug/mL Puromycin (Sigma, P8833) was added to SK-N-AS and GI-ME-N cells respectively. Several weeks after transfections SK-N-AS cells were single-cell sorted by FACS using the SH800S (Sony Biotechnology) sorter in 96-well plates and GI-ME-N cells were single-cell diluted in 96-well plates. Clonal cultures were expanded and harvested for gDNA, protein and RNA to confirm editing.

*ATRX* MED clones were established by transfecting GI-ME-N and NB139 cells with two Cas9 expressing vectors containing two distinct sgRNAs (Table S4) using Fugene HD transfection reagent. Three days after transfection, GFP-positive (transiently expressed) cells were single-cell sorted and expanded as described above. Genomic DNA, protein and RNA was harvested to confirm genome editing. For GI-ME-N, we expected only one *ATRX* allele (X-chromosomal loss according to WGS data), however our GI-ME-N cells contained three alleles. Presence of MEDs were confirmed on only one allele in all GI-ME-N clones. Subsequently, we transfected GI-ME-N cells with an *ATRX* exon 2-13 MED with ATRX_KO_sgRNA_1 and the corresponding homology arm plasmid to KO the remaining wild-type alleles using Fugene HD transfection reagent. While for some GI-ME-N clones with an exon 2-10 MED, we transfected them with either ATRX_KO_sgRNA_1 (clone 7, 31 and 39) or ATRX_KO_sgRNA_2 (all other clones) and the corresponding homology arm plasmids using Fugene HD transfection reagent. Three days after transfection, medium containing 1 μg /mL Puromycin was added to the cells and several days or weeks later, cells were single-cell sorted by FACS in 96-well plates. Clonal cultures were expanded and harvested for gDNA, protein and RNA to confirm editing.

CHLA-90 doxycycline-inducible *ATRX* wild-type clones were established by transfecting cells with 50 ng PiggyBac doxycycline-inducible mCMV-Kozak-ATRX-wildtype plasmid and 50 ng of the pCMV-hyPBase plasmid (a gift from Kosuke Yusa^44^, Wellcome Trust Sanger Institute) in 12-well plates. Three days after transfection, medium containing 1000 μg/mL Neomycin (G-418, Roche) was added to the cells. Several weeks later, cells were single-cell sorted by FACS in 96-well plates. Clonal cultures were expanded, and protein was harvested for cells treated with and without 2500 ng/ml doxycycline to confirm editing.

### gDNA and cDNA Validation of clones

For genotyping, we extracted genomic DNA utilizing the Wizard® SV Genomic DNA Purification System (Promega). Primers were designed to amplify the allele with the PGK-eGFP-Puromycin insert, the wild-type allele or the allele containing distinct MEDs (Table S6). For mRNA expression validations, cDNA was generated using 2.5 microgram of RNA and the IScript cDNA Synthesis Kit (1708891, Bio-Rad) according to the manufacturer’s manual. Primers were designed to amplify the wild-type or the distinct MEDs allele mRNA products. PCR products were analysed by gel electrophoresis and Sanger sequencing.

### Western blot analysis

Western blots were performed as described in Boogaard et al^42^. Primary and secondary antibodies are shown in Table S7.

### ALT Southern blot analysis

DNA for southern blot was isolated in 10 ml SE buffer (75mM NaCl, 25mM Na2, EDTA, pH 8.0) and extracted using phenol-chloroform extraction. Southern blots to detect ALT were performed using the TeloTAGGG Telomere Length Assay Kit (12209136001, Sigma) according to the manufacturers protocol.

### ALT-associated PML Bodies (APBs) staining and immunofluorescent (IF) staining

APBs and IF stainings were performed on cells grown on coverslips in 6-well plates. For APBs stainings, coverslips were washed twice with 1x PBS and incubated with 4% paraformaldehyde for 20 minutes. Subsequently, cover glasses were incubated for 3 minutes with 70%, 2 minutes with 95% and 2 minutes with 100% ethanol. Next, coverslips were airdried and put on a coverglass with 10 μL of probe in-between (10 μL HB buffer (50% formamide, 10% Dextran Sulfate Sodium and 2x SSC (saline-sodium citrate)) and 0.5 μL TelC-Alexa-Fluor-488 PNA Bio probe (F1104)). Subsequently, the coverslips were fixed on the coverglass using Fixogum and airdried for 1 hour. Next, denaturation was performed for 2 minutes at 75 °C and the slides were incubated overnight in a wet hybridization chamber at 37 °C. The next day, the Fixogum and coverglass were removed and coverslips were washed with 2x SSC buffer for 5 minutes, followed by 1x PBS wash. Subsequently, permeabilization was performed for 5 minutes using 1x PBS/0.1X triton X-100 followed by 1x PBS wash. Next, coverslips were incubated for 5 minutes with 10 mM Sodium Citrate (pH 6), after which 30 minutes blocking was performed using TBS/1% BSA/0.1% Triton X-100. Subsequently, coverslips were incubated overnight at 4 °C with the primary PML antibody (See Table S7). The next day, coverslips were washed three times with 1x PBS and incubated with the secondary antibody (Table S7) for 2 hours at room. Subsequently, coverslips were washed 3 times with 1x PBS followed by incubation for 3 minutes with 70%, 2 minutes with 95% and 2 minutes with 100% ethanol. Lastly, coverslips were airdried, DAPI stained and sealed on a coverglass.

For IF stainings, coverslips were washed twice with 1x PBS/0.1% Tween-20 (PBS-T) and cells were fixed for 20 minutes using 4% paraformaldehyde. Subsequently, coverslips were washed 3 times with 1x PBS-T and permeabilization was performed using 1x PBS/0.2% Tween-20. Next, coverslips were washed 3 times with 1x PBS-T, after which 30 minutes blocking with 1% BSA in PBS-T was conducted. Subsequently, coverslips were incubated overnight at 4 °C with the primary antibodies (Table S7). The next day, coverslips were washed three times with 1x PBS-T and incubated with the secondary antibodies (Table S7) for 1 hours at room temperature. Subsequently, coverslips were washed 3 times with 1x PBS-T and coverslips were airdried, DAPI stained and sealed on a coverglass. Imaging for both APBs and IF stainings was performed on a Leica DM RA microscope with a 63 mm lens.

### Cell cycle and Ki67 analysis

Cells were harvested, washed in 1x PBS and resuspended in 1x PBS 2mM EDTA until single cells were obtained. Subsequently, cells were washed with 1x PBS and 1 million cells were stained with 1x PBS containing 1:1000 Zombie NIR™ (BioLegends). Cells were incubated for 20 minutes at room temperature and washed with 1x PBS. Subsequently, cells were fixated with 200 μL fixation buffer (eBioscience™ Foxp3/Transcription Factor Staining Buffer, Set, Invitrogen™) and incubated for 30 minutes at 4 °C. Next, cells were washed twice with 500 μL permeabilization buffer (from kit described above) and stained with 150 μL permeabilization buffer containing 1:400 Ki67-Alexa-Fluor antibody (561126, BD Biosciences) for 30 minutes at 4 °C. Subsequently, cells were washed with permeabilization buffer and FACS buffer (2% FCS 2mM EDTA 1x PBS), after which the cells were stained in FACS buffer containing 1:500 Vybrant® DyeCycle™ Green (V35005, Invitrogen) for 30 minutes at 37 °C. Next, FACS was performed on a CytoFLEX S flow cytometer (Beckman Coulter) and analyses were conducted using CytExpert and FloJo software.

### Violet trace analysis

Cells were washed with 1x PBS and stained with 1x PBS containing 2 μM CellTrace™ Violet (C34557, Invitrogen) for 7 minutes at 37 °C. Subsequently, 10 volumes ice-cold FBS were added and the cell mixture was centrifuged for 10 minutes at 250 g at 4 °C. Next, cells were washed twice with medium and plated. Two days later, cells were harvested and stained with Zombie NIR™ and resuspended in FACS buffer for FACS analysis as described above.

### RNA extraction and purification for RNA sequencing and qPCRs

Cells were harvested in TRIzol™ (Invitrogen), and chloroform extraction was performed. RNA was precipitated using 100% RNA-free ethanol to the aqueous phase and eluted in 100 uL MilliQ. Subsequently, the RNA was further purified using the NucleoSpin RNA kit (Macherey-Nagel) according to manufacturer’s protocol.

### rRNA qRT-PCRs and analysis

Five microgram of RNA were used for cDNA synthesis using random primers and the SuperScript™ II Reverse Transcriptase kit (18064014, Invitrogen) according to the manufacturer’s protocol. qRT-PCRs were performed on a C1000 thermal cycler (Bio-Rad) using SYBR green (170886, Bio-Rad) for the primers listed in Table S8. For each primer set, triplicate reactions were performed and data analysis was performed by using the CFX Maestro Software (Bio-Rad). Expression levels were normalized to ATP5PO and UBE3B expression. Biological replicates were averaged and normalized against wild-type average minus 1.

### RNA sequencing and analysis

For our isogenic model systems, we wanted to compare at least four mutant clones versus four wild-type clones. However, not for all isogenic models we acquired four clones (Table S1) and therefore we split the cells to the required number of biological replicates. Before harvesting, we first maintained the splits of cells separately for at least 1.5 week, so that new independent mutations could be acquired.

For all isogenic clones, Illumina sequencing libraries were prepared using the Truseq RNA stranded RiboZeroPlus Kit (Illumina) and sequenced with 2×50bp paired-end sequencing on an Illumina Novaseq S1 (1500M) System. After sequencing the data was aligned to genome build GRCh37 (gencode v74) using STAR(v2.7.3a) and counts were generated using the R package Rsubread^45^. For all cell lines, Illumina sequencing libraries were prepared using the KAPA RNA HyperPrep Kit with RiboErase Kit (Roche) and sequenced with 2×150bp paired-end sequencing on an Illumina NovaSeq6000 System. After sequencing the data was aligned to genome build GRCh38 (gencode v19) using STAR(version 2.7.0f) and counts were generated using the R package Rsubread^45^. For all iTHER tumour, Illumina sequencing libraries were prepared using the TruSeq RNA V2 Kit (Illumina) and sequenced with 2×100bp paired-end sequencing on an Illumina HiSeq4000 System. After sequencing the data was aligned to genome build GRCh37 (gencode v17) using STAR(version 2.3.0e) and counts were generated using the R package Rsubread^45^.

For all three datasets, counts were normalized by Variance Stabilizing Transformation (VST from DESeq2 package) and differentially expressed genes with a adjusted p-value of <0.05 were determined using the DESeq2^46^ R package. For the GI-ME-N clones, we observe a batch effect in the wild-type clones. Therefore, we decided to compare the different GI-ME-N *ATRX* aberrant models only with their corresponding wild-type clones (generated by same person). Gene ontology analysis was performed by inputting the overlapping differentially expressed genes in the gene ontology resource (http://geneontology.org/) for gene ontology biological process and Panther protein class datasets using Fisher exact test and FDR correction. GSEA was conducted using the R package fgsea with the gene ontology biological process and reactome gene set databases from MsigDB V7.2^47^. Heatmaps were generated using the R package pheatmap^48^, proportional Venn diagrams were created utilizing the eulerr^49,50^ package and the remaining figures were generated using the package ggplot2^51^.

## Acknowledgements

The research in this paper was supported by funding from the European Research Council (ERC) under the European Union’s Horizon 2020 research and innovation programme under grant agreement numbers 716079 (Predict) and 826121 (iPC project).

