## Supplements for "Two opposing gene expression patterns within *ATRX* aberrant neuroblastoma"

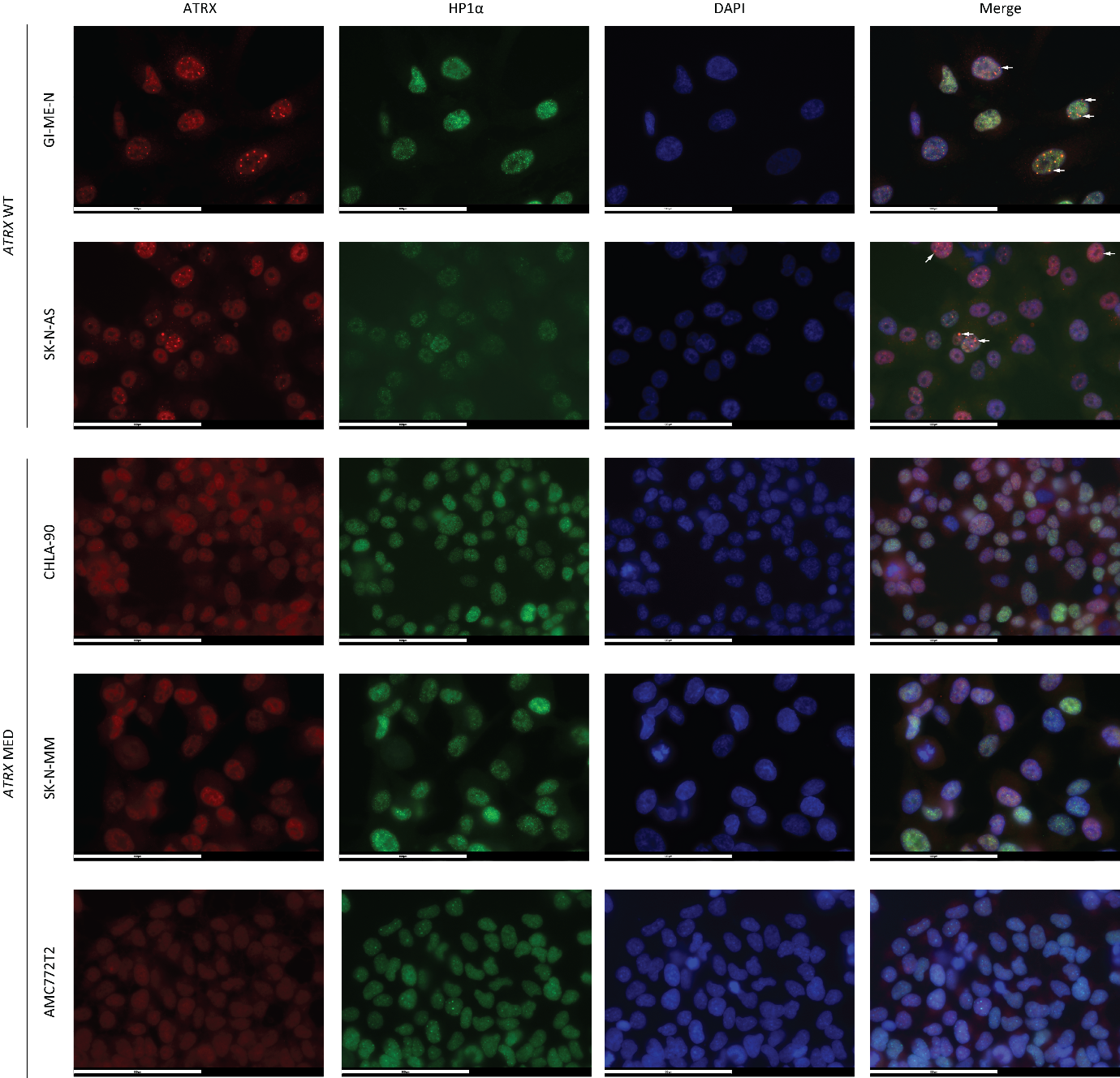

**Supplementary figure 1. ATRX immune-fluorescent staining confirms diffuse ATRX staining in three patient-derived *ATRX* MED models and absence of HP1α co-localisation.** White arrows mark co-localisation of ATRX and HP1α foci, only a maximum of four arrows per image is displayed.

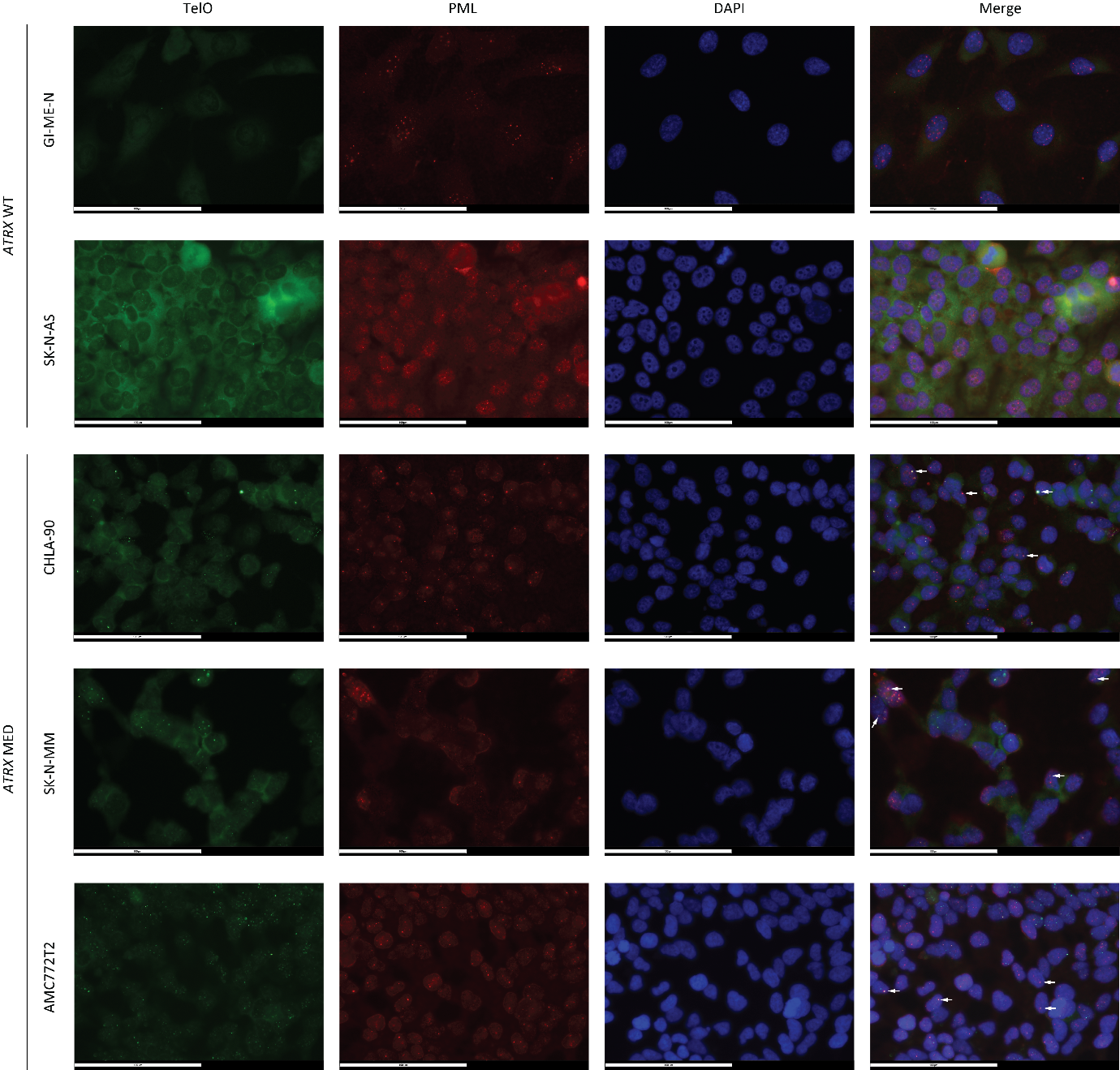

**Supplementary figure 2. ALT-associated PML bodies (APBs) staining confirms the present of ALT in three patient-derived *ATRX* MED models.** White arrows mark co-localisation of telomeric (TelO) and PML foci, only a maximum of four arrows per image is displayed.

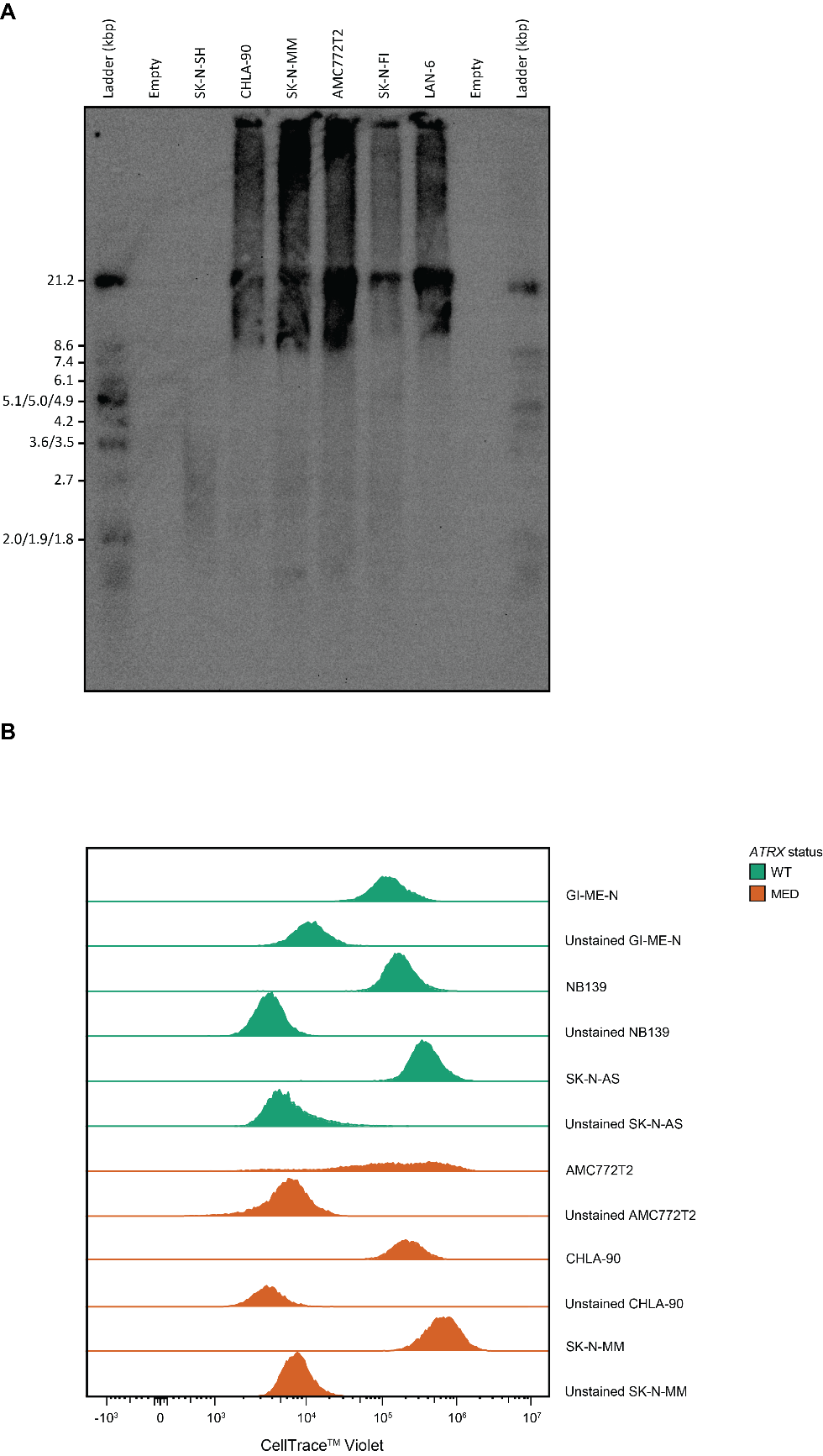

**Supplementary figure 3. Southern blot confirms the present of ALT in three patient-derived *ATRX* MED models and violet trace identified unchanged proliferation rates. A)** Southern blot containing one ALT negative cell line (SK-N-SH) and two well-known ALT positive neuroblastoma cell lines (SK-N-FI and LAN-6). All three PD^ΔATRX^ models display long and heterogeneous telomeres and therefore confirm ALT. **B)** Violet trace experiments on three *ATRX*^WT^ and on three PD^ΔATRX^ models.

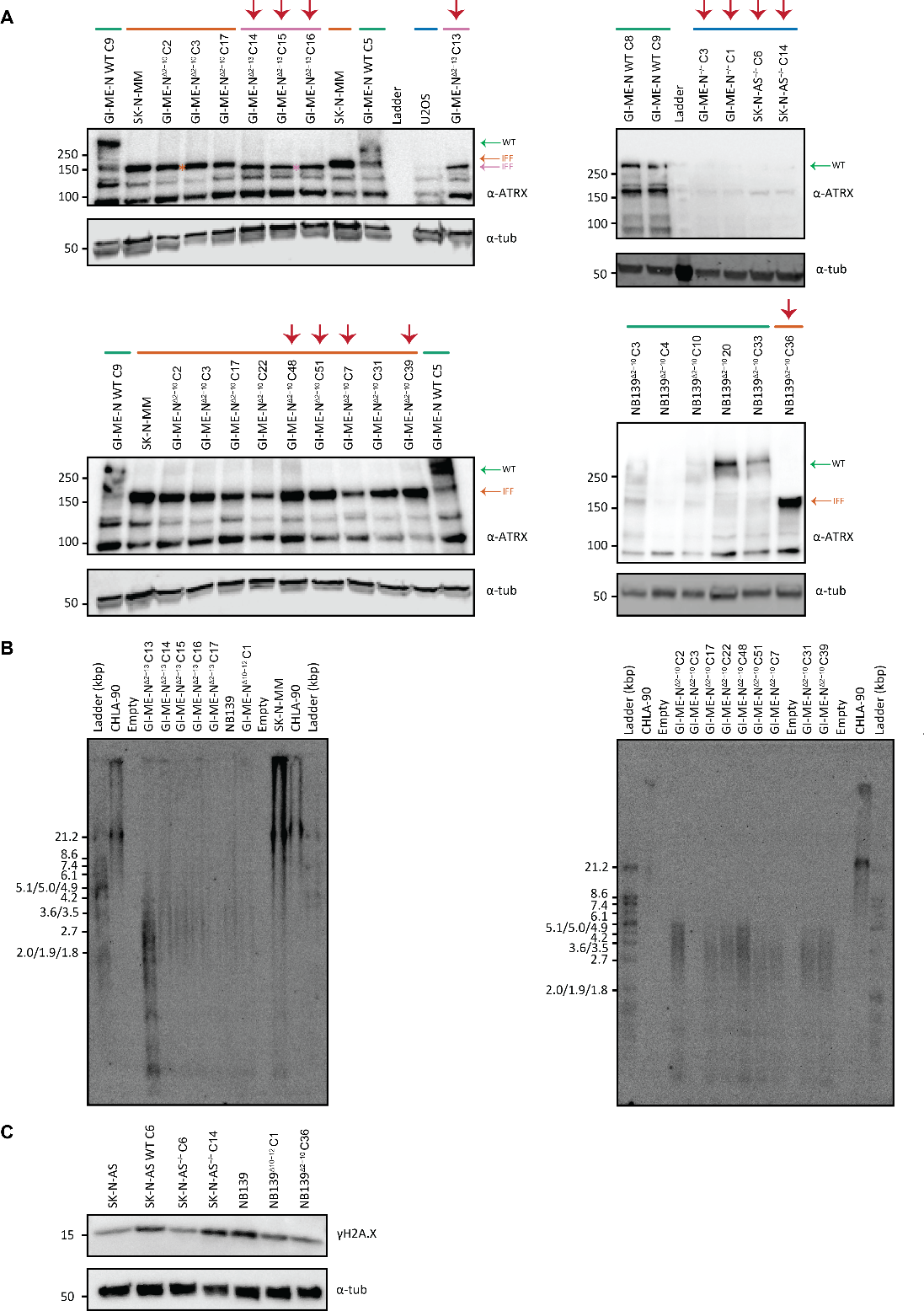

**Supplementary figure 4. Western blot confirmation of the generated *ATRX* aberrant isogenic models and absence of ALT. A)** Western blots for ATRX protein of all the correct *ATRX* aberrant clones. Clones that were send for sequencing are marked with a red arrow on top. Green bar: confirmed wild-type clones, dark orange bar: confirmed *ATRX*^Δ2-10^ clones or PD^ΔATRX^ models, pink bar: confirmed *ATRX*^Δ2-13^ clones and dark blue bar: confirmed *ATRX*^-/-^ clones or the *ATRX*^-/-^ osteosarcoma cell line U2OS. IFF: *ATRX* in-frame fusion protein product. **B)** Southern blots confirming absence of long heterogeneous telomeres in our isogenic *ATRX* aberrant models. CHLA-90 and SK-N-MM were used as positive controls. C) Western blot of γH2A.X abundance in isogenic *ATRX* aberrant SK-N-AS and NB139 models compared to wild-type motherlines and clone.

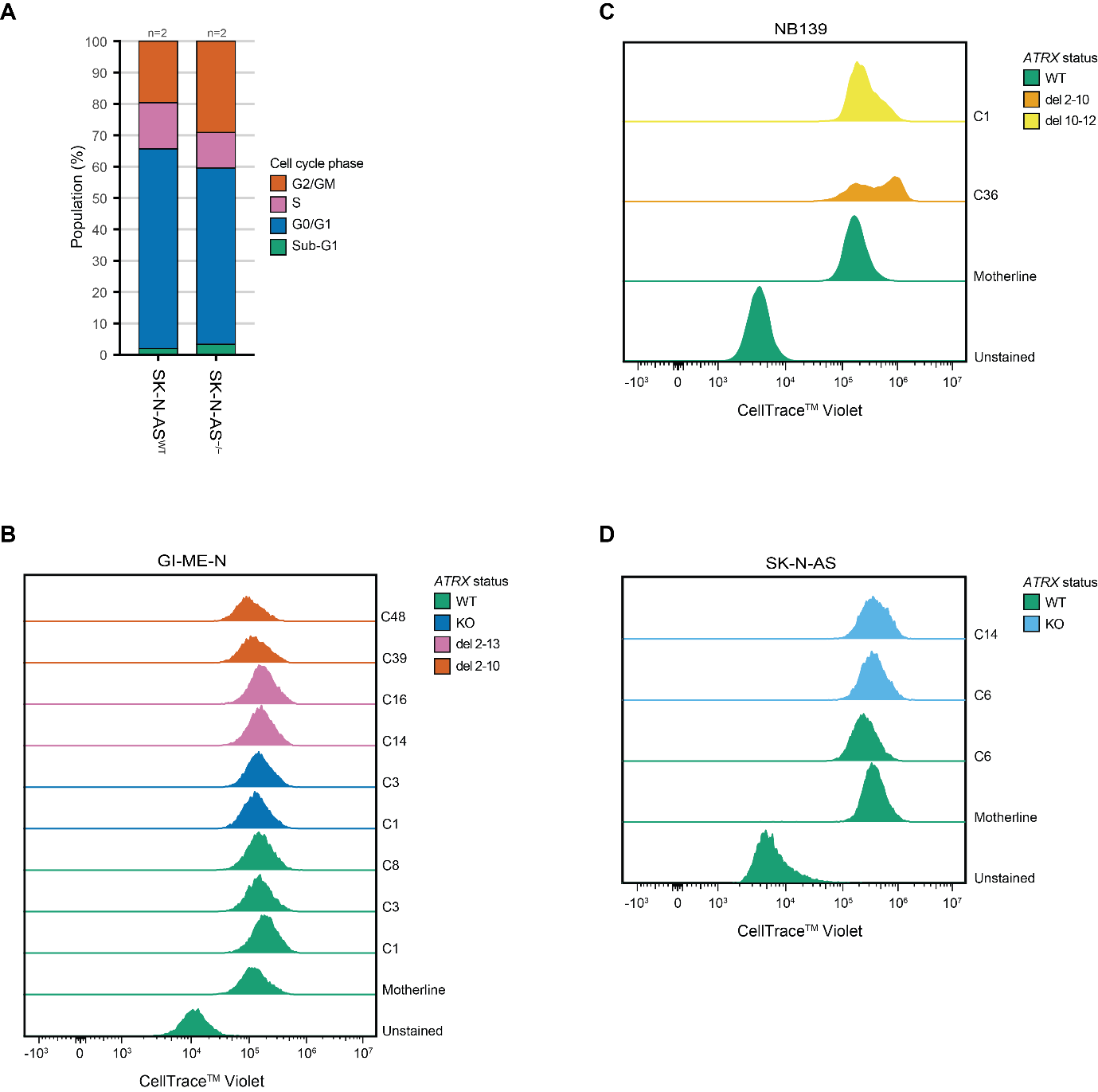

**Supplementary figure 5. Unchanged cell cycle progression and proliferation in SK-N-AS and unaltered proliferation rates in the majority of the isogenic *ATRX* aberrant models. A)** Cell cycle distribution analysis of *ATRX* wild-type and isogenic *ATRX* aberrant SK-N-AS models. “n=” indicates the number of biological replicates used in these experiments. **B-D)** Violet trace experiments on *ATRX* wild-type and isogenic *ATRX* aberrant **B)** GI-ME-N, **C)** NB139 and **D)** SK-N-AS models.

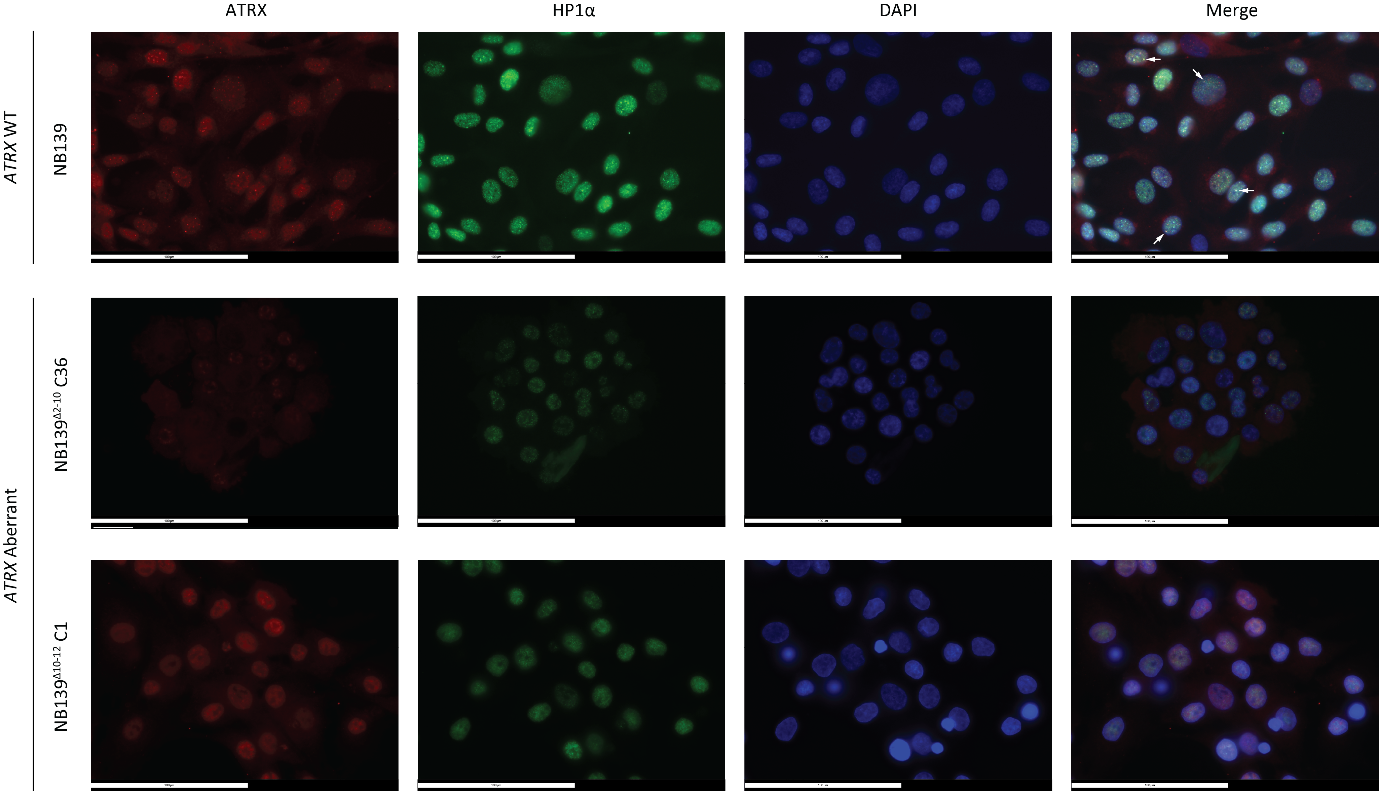

**Supplementary figure 6. ATRX immune-fluorescent staining confirms diffuse ATRX staining in isogenic *ATRX* aberrant NB139 models and absence of HP1α co-localisation.** White arrows mark co-localisation of ATRX and HP1α foci, only a maximum of four arrows per image is displayed.

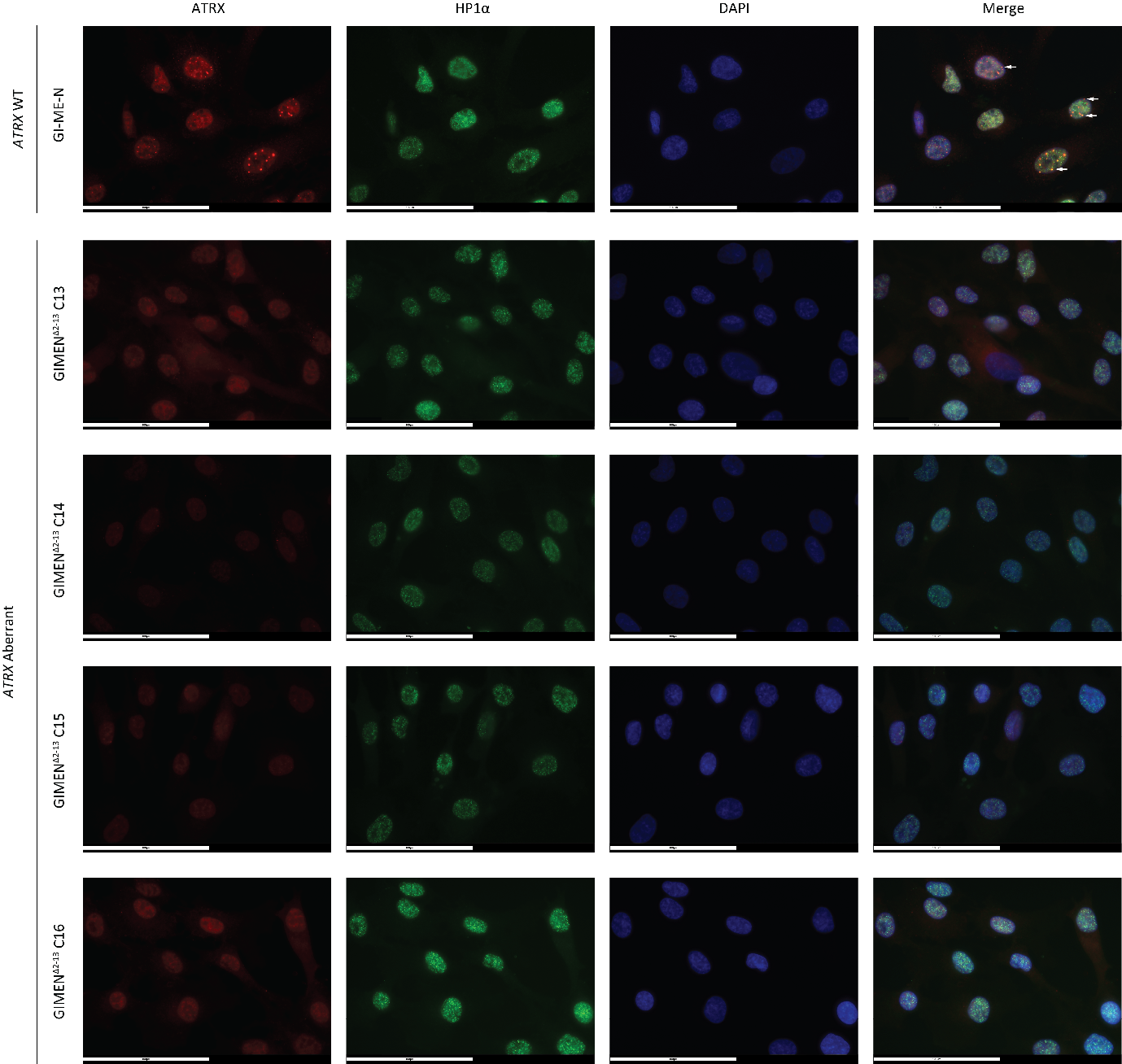

**Supplementary figure 7. ATRX immune-fluorescent staining confirms diffuse ATRX staining in isogenic *ATRX* aberrant GI-ME-N^Δ2-13^ models and absence of HP1α co-localisation.** White arrows mark co-localisation of ATRX and HP1α foci, only a maximum of four arrows per image is displayed.

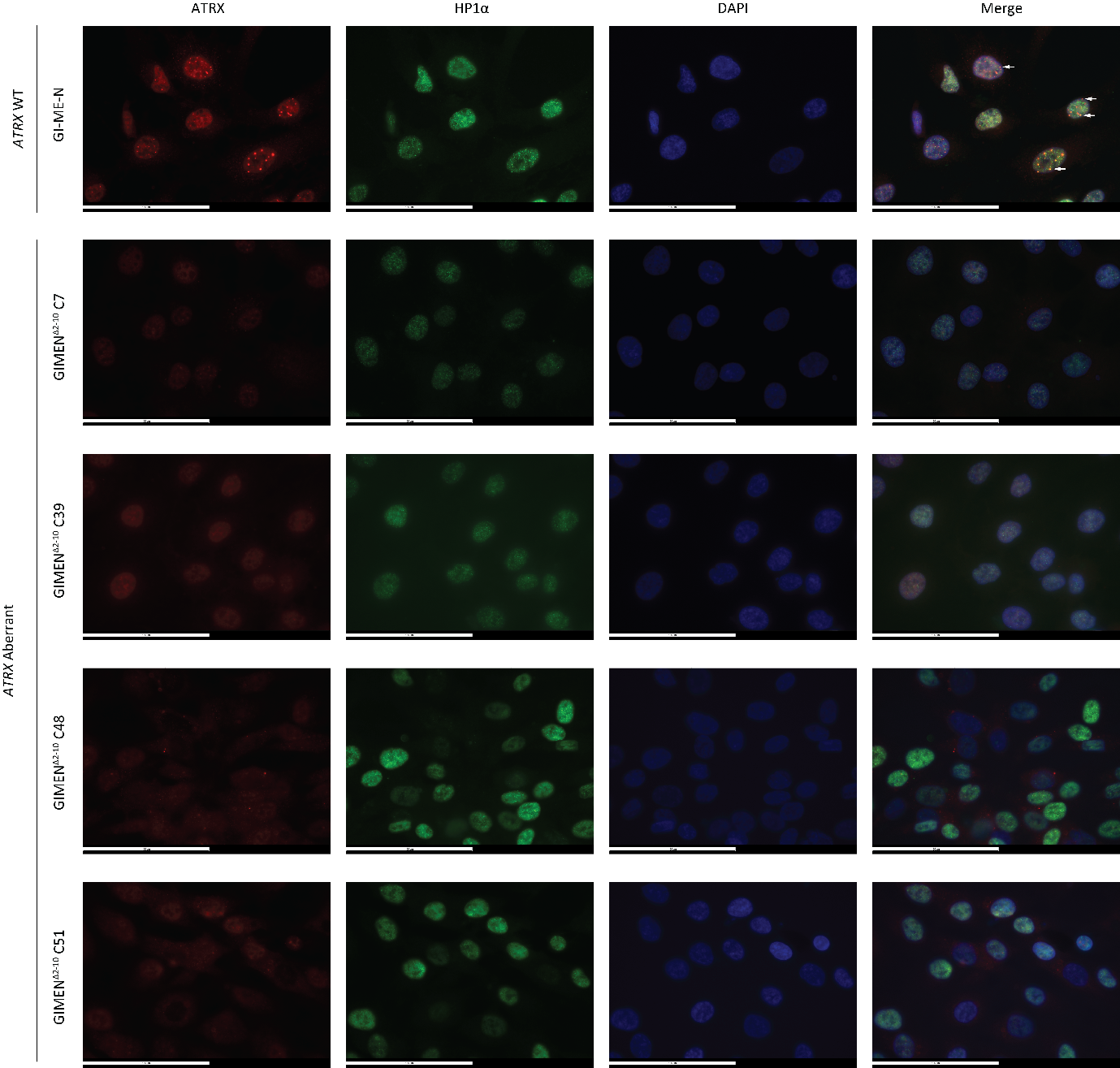

**Supplementary figure 8. ATRX immune-fluorescent staining confirms diffuse ATRX staining in isogenic *ATRX* aberrant GI-ME-N^Δ2-10^ models and absence of HP1α co-localisation.** White arrows mark co-localisation of ATRX and HP1α foci, only a maximum of four arrows per image is displayed.

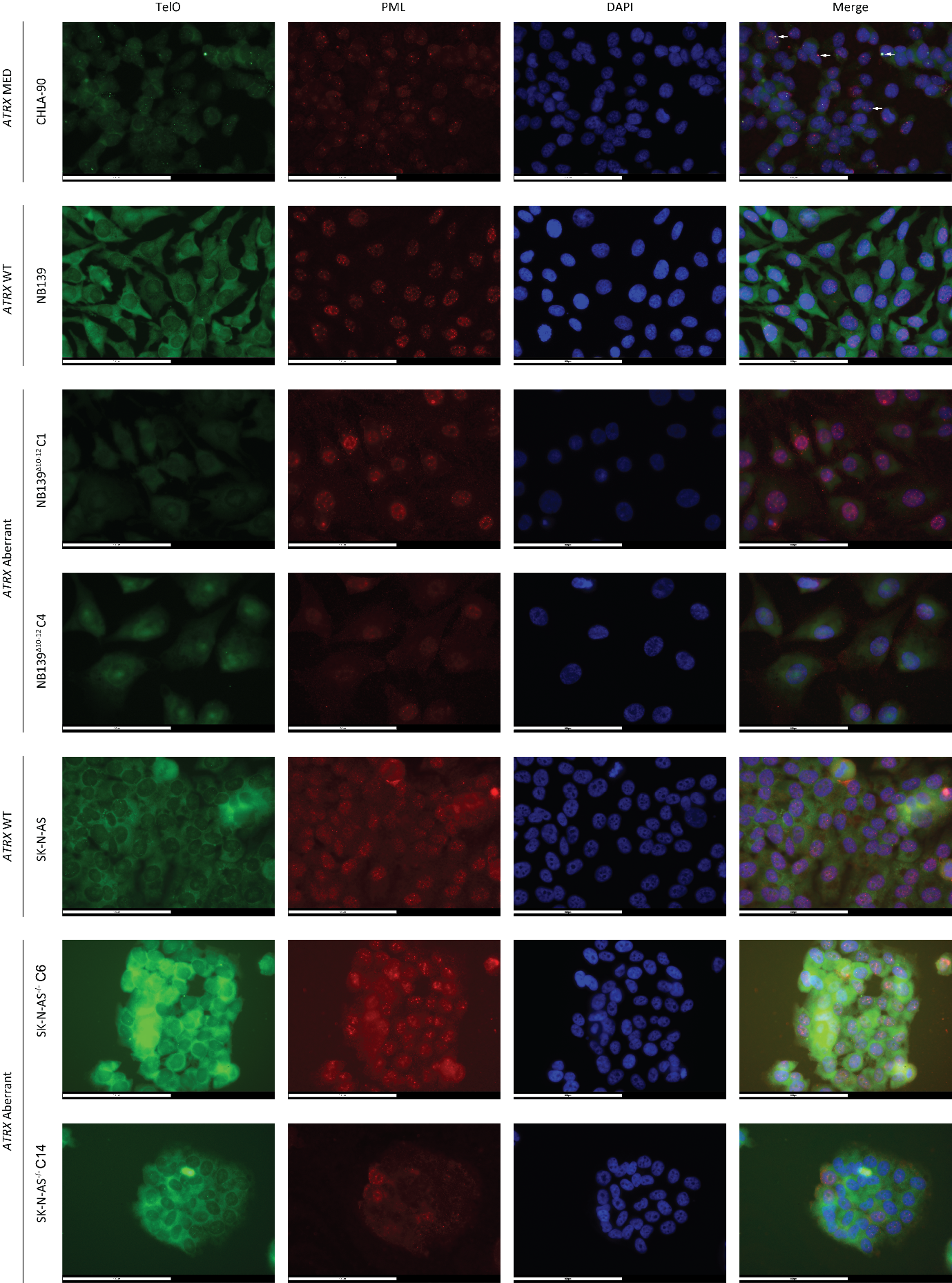

**Supplementary figure 9. ALT-associated PML bodies (APBs) staining identifies absence of ALT in isogenic *ATRX* aberrant NB139 and SK-N-AS models.** White arrows mark co-localisation of telomeric (TelO) and PML foci, only a maximum of four arrows per image is displayed. CHLA-90 was added as positive control.

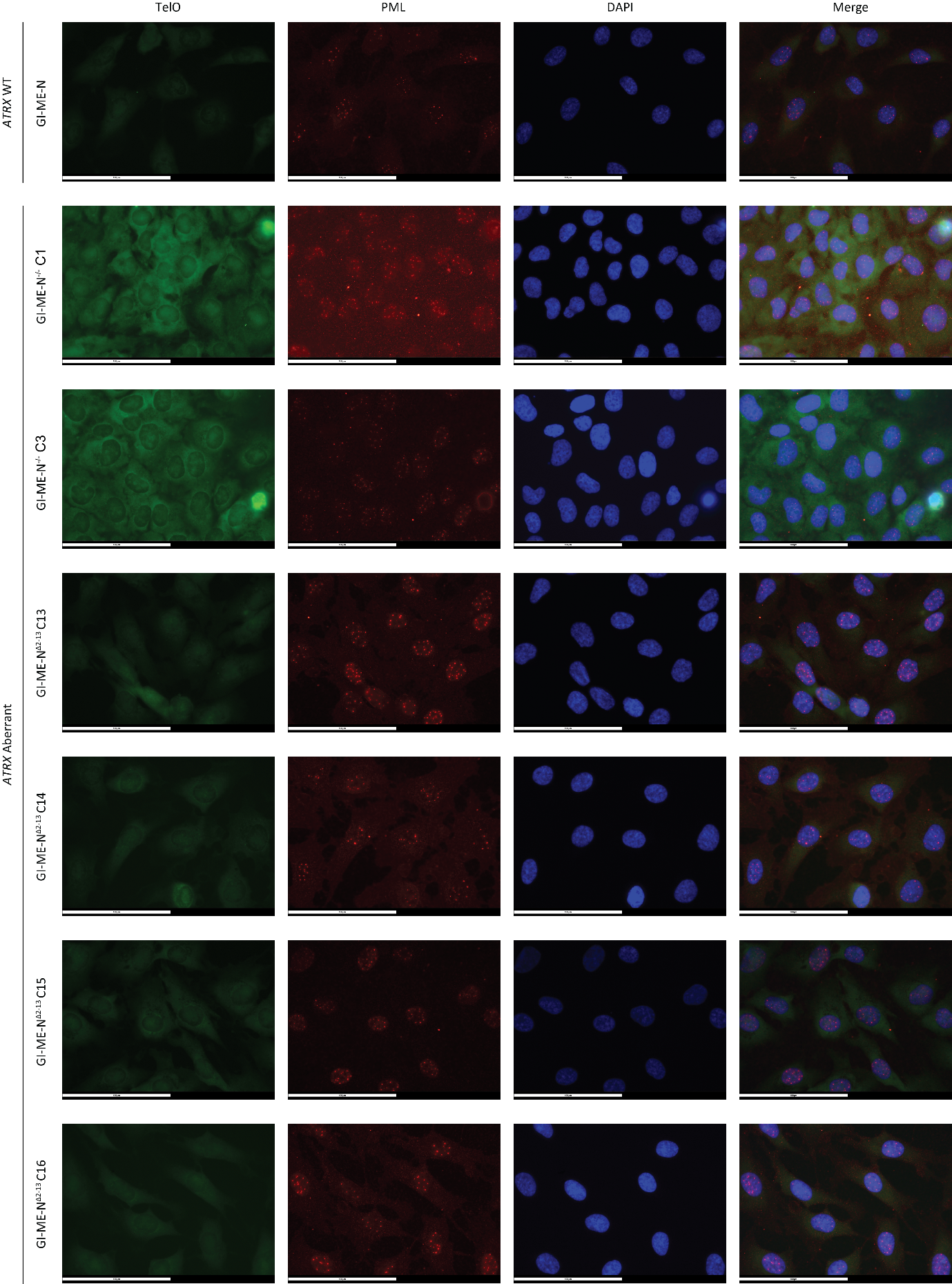

**Supplementary figure 10. ALT-associated PML bodies (APBs) staining identifies absence of ALT in isogenic *ATRX* aberrant GI-ME-N^-/-^ and GI-ME-N^Δ2-13^ models.**

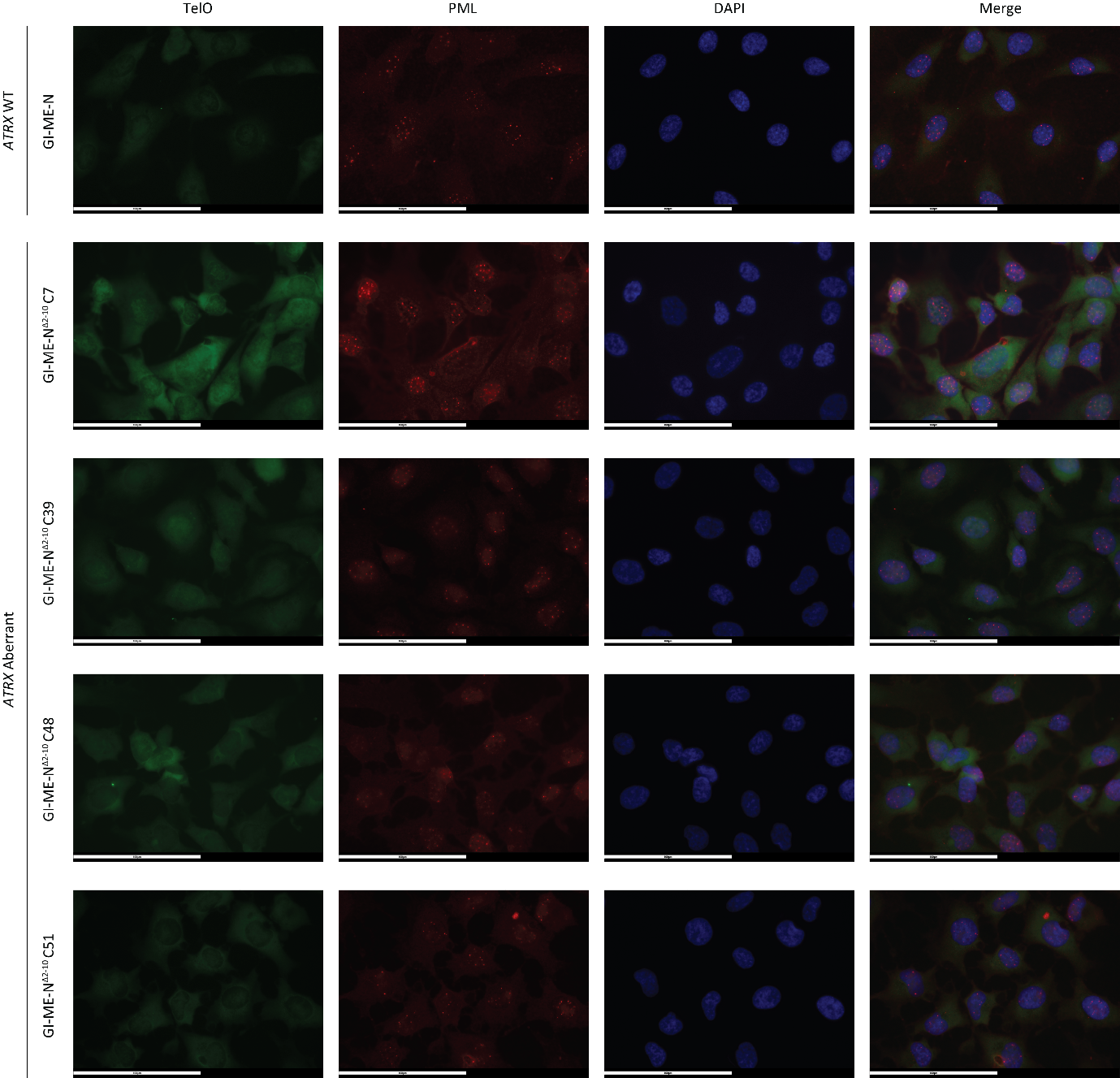

**Supplementary figure 11. ALT-associated PML bodies (APBs) staining identifies absence of ALT in isogenic *ATRX* aberrant GI-ME-N^Δ2-10^ models.**

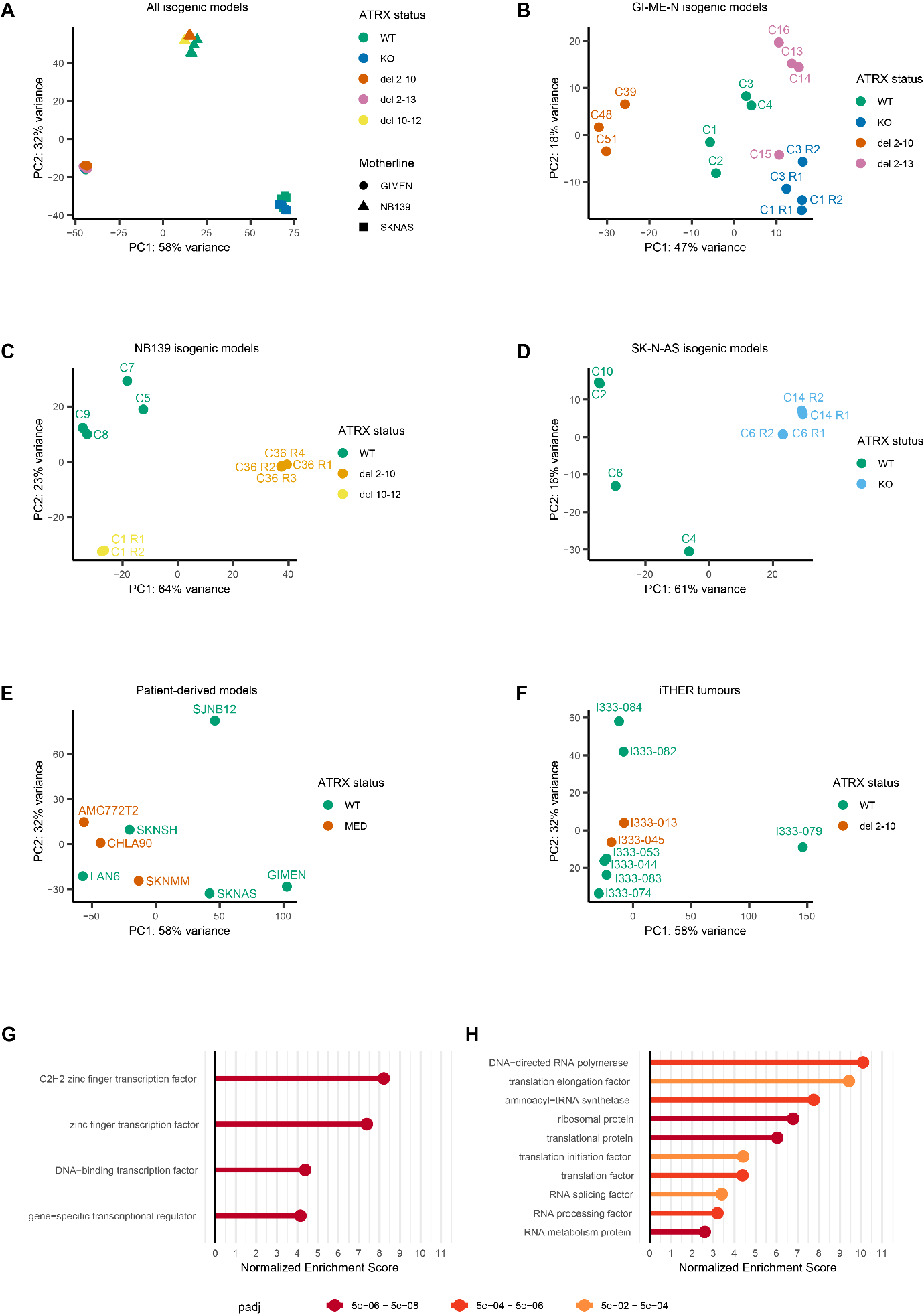

**Supplementary figure 12. Principle component analysis (PCA) and Panther protein class analysis of generated isogenic *ATRX* aberrant models and PCA of patient-derived models and iTHER tumours. A)** PCA of all generated isogenic *ATRX* aberrant clones showing separation based on the motherliness. **B-F)** PCA of **B)** isogenic GI-ME-N clones, **C)** isogenic NB139 clones, **D**) isogenic SK-N-AS clones, **E)** patient-derived *ATRX* aberrant and wild-type models **F)** *ATRX* aberrant and wild-type iTHER tumours. **G-H)** Significantly enriched Panther protein classes of the overlapping differentially expressed downregulated genes **G)** for all *ATRX^-/-^* and *ATRX*^Δ2-13^ isogenic models and **H)** for all *ATRX*^Δ2-10^ isogenic models.

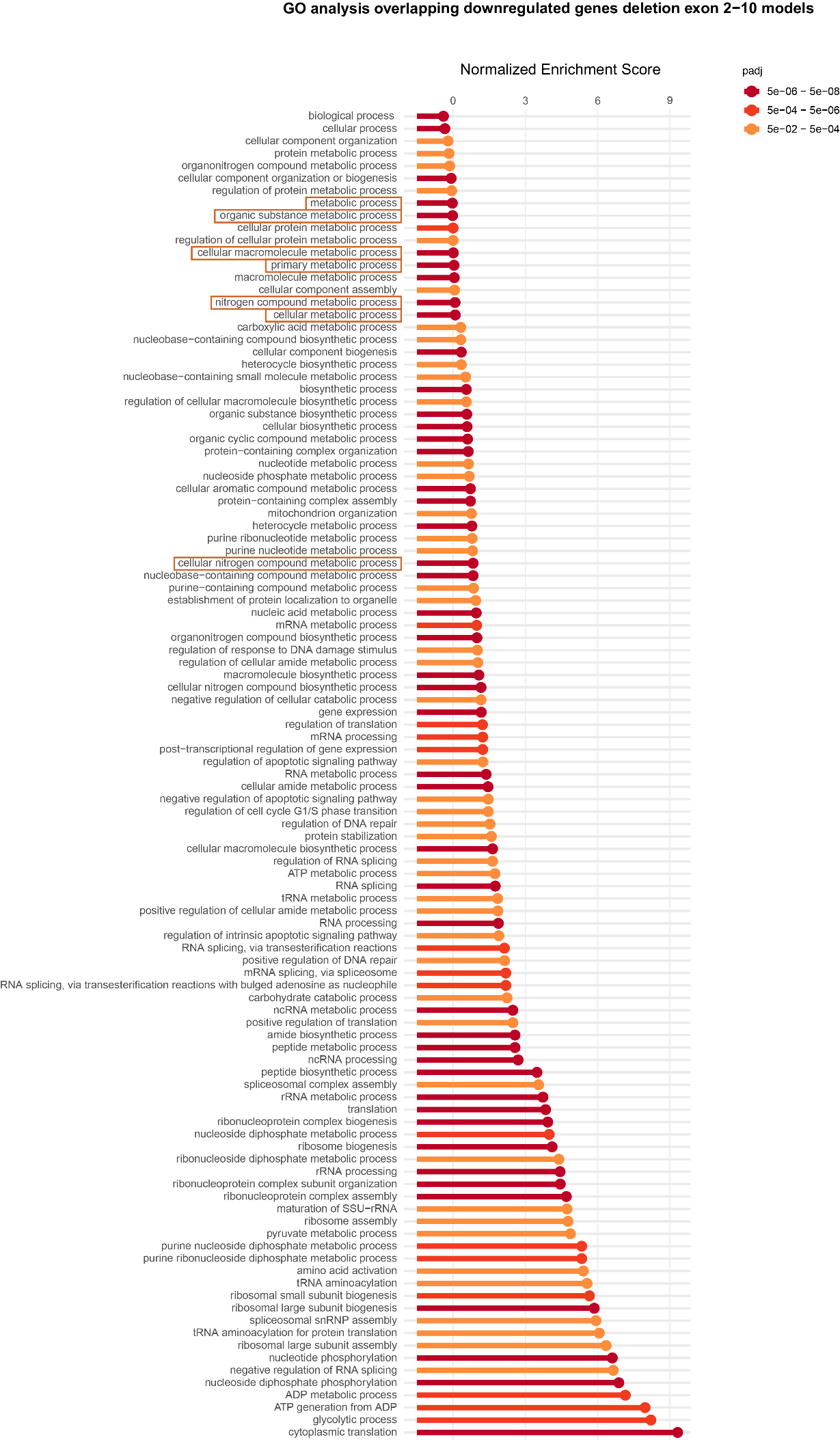

**Supplementary figure 13. The overlapping differentially expressed downregulated genes of all ATRX^Δ2-10^ isogenic models are enriched for distinct RNA and metabolic processes according to GO BP.** Orange boxes highlighted the same terms as observed in figure 3C.

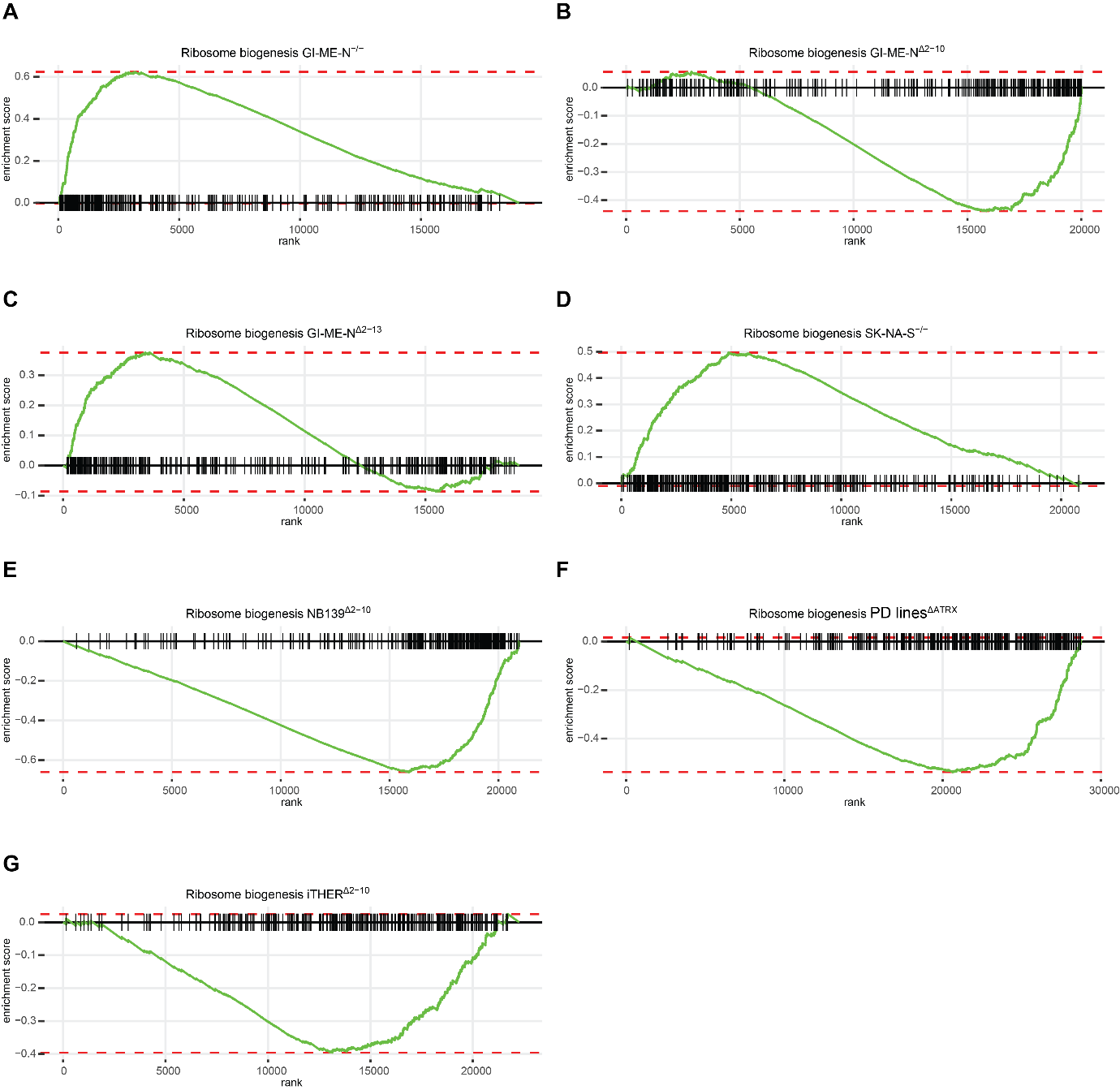

**Supplementary figure 15. Enrichment plots for all our performed GSEA for the GO BP gene set ribosome biogenesis.**

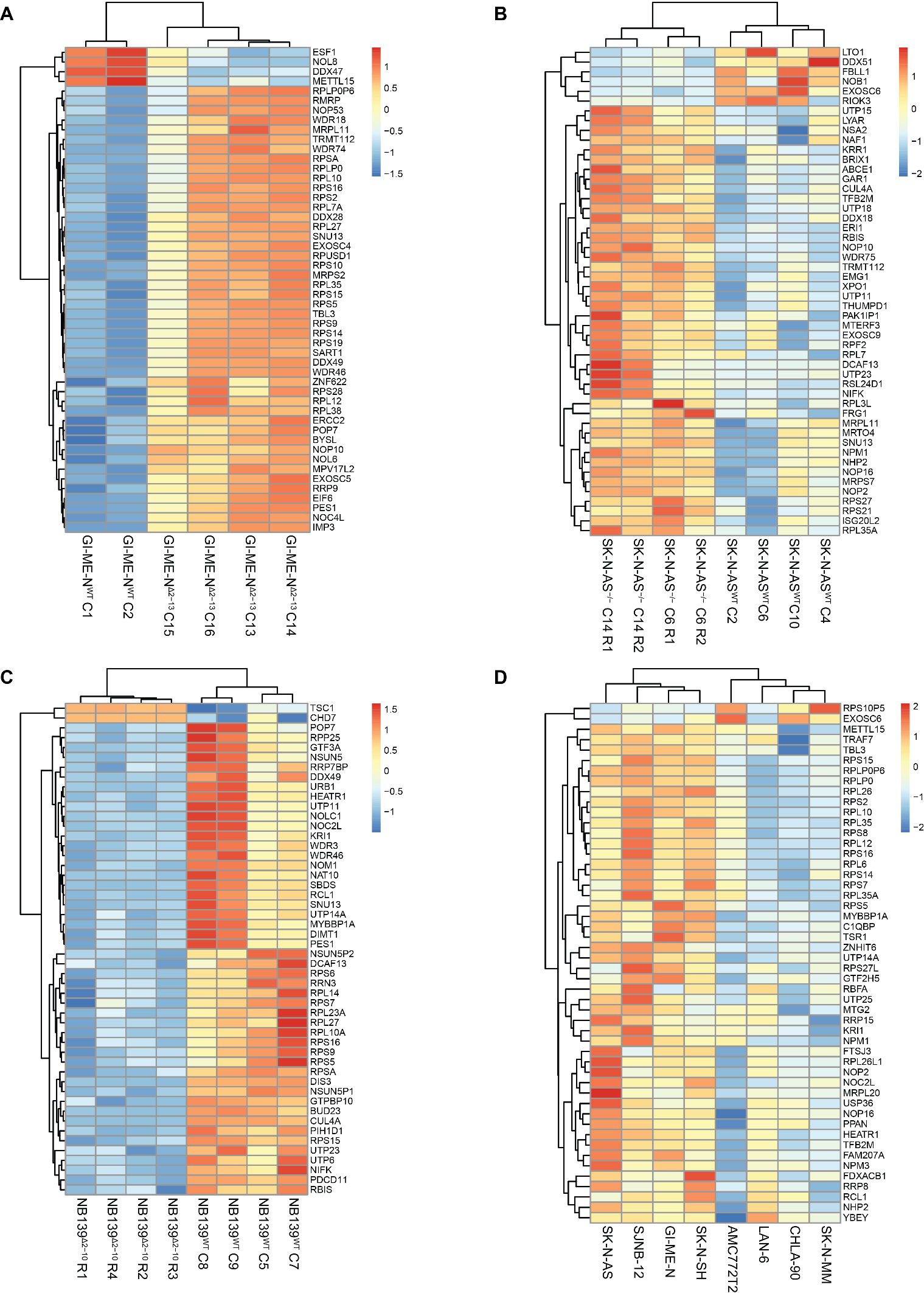

**Supplementary figure 15. Gene expression of the top 50 differentially expressed ribosome biogenesis genes confirms changes in ribosome biogenesis in *ATRX* isogenic and patient-derived models. A-D)** Heatmaps showing expression values for the top 50 differentially expressed ribosome biogenesis genes that are normalized across all samples by Z-score. Both row and column clustering were applied, and distinct clusters were identified.

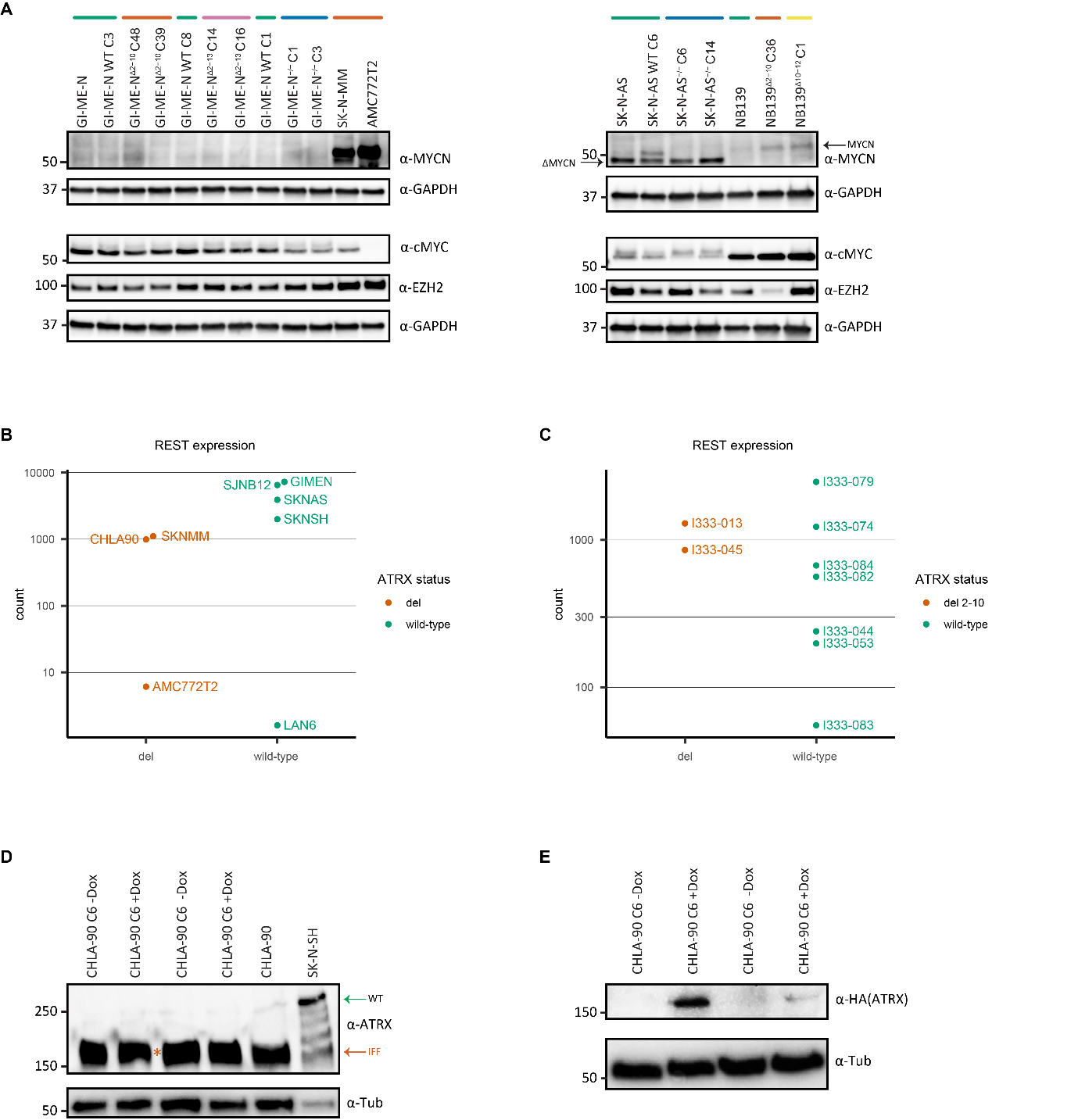

**Supplementary figure 16. Unchanged gene expression and protein expression in the majority of our isogenic models for several proteins that are involved in ribosome biogenesis. A)** Western blot analysis for three proteins (MYCN, cMYC and EZH2) that are directly involved in regulating ribosome biogenesis reveals unchanged expression in the majority of our isogenic models and no changes that are consistent with our identified expression pattern dichotomy within *ATRX* aberrant models. Two distinct isoforms of MYCN are displayed in the right blot, with SK-N-AS being the only model expressing ΔMYCN **B)** REST gene expression in patient-derived *ATRX* MED and wild-type models. **C)** REST gene expression in *ATRX* aberrant and wild-type iTHER tumours. **D)** Western blot analysis showing unchanged *ATRX* IFF protein expression upon doxycycline induction of ATRX^WT^ protein expression. **E)** Western blot analysis showing presence of HA-tagged ATRX^WT^ protein product only upon induction with doxycycline.

**Supplementary figure 1. Genetic overview of (un)successful clones validated by Sanger sequencing on both genomic DNA and RNA (cDNA).** RNA confirmed ‘Yes’ means absence of wildtype mRNA and for the MED models also detection of mRNA expression of the deleted allele. X-Chr: X-Chromosome.

| Model/clone name | Success-  ful | Female  /Male | Number of X-Chr | Allele 1 | Allele 2 | Allele 3 | Passage cells | RNA confirmed |
| --- | --- | --- | --- | --- | --- | --- | --- | --- |
| SK-N-AS^-/-^ C6 | Yes | F | 2 | Insert A exon 4 | Insert A exon 4 | - | 1:5 | Yes |
| SK-N-AS^-/-^ C14 | Yes | F | 2 | Insert A exon 4 | Insert A exon 4 | - | 1:3 | Yes |
| GI-ME-N^-/-^ C1 | Yes | F | 3 | PuroR | del exon 4 (del 87bp and 5bp inserted) | Insert A exon 4 | 1:10 | Yes |
| GI-ME-N^-/-^ C3 | Yes | F | 3 | PuroR | del exon 4 (del 516bp and 1bp inserted) | del exon 4 (del 453bp) | 1:10 | Yes |
| GI-ME-N^Δ2-10^ C2 | Yes | F | 3 | PuroR | PuroR | del ex2-10 | 1:10 | Yes |
| GI-ME-N^Δ2-10^ C3 | Yes | F | 3 | PuroR | PuroR | del ex2-10 | 1:10 | Yes |
| GI-ME-N^Δ2-10^ C7 | Yes | F | 3 | PuroR | Del 2nt exon 4 | del ex2-10 | 1:10 | Yes |
| GI-ME-N^Δ2-10^ C17 | Yes | F | 3 | PuroR | PuroR | del ex2-10 | 1:10 | Yes |
| GI-ME-N^Δ2-10^ C22 | Yes | F | 3 | PuroR | Insert 1nt exon 4 | del ex2-10 | 1:10 | Yes |
| GI-ME-N^Δ2-10^ C31 | Yes | F | 3 | PuroR | del 38nt exon 4 | del ex2-10 | 1:10 | Yes |
| GI-ME-N^Δ2-10^ C39 | Yes | F | 3 | PuroR | PuroR | del ex2-10 | 1:10 | Yes |
| GI-ME-N^Δ2-10^ C48 | Yes | F | 3 | PuroR | PuroR | del ex2-10 | 1:10 | Yes |
| GI-ME-N^Δ2-10^ C51 | Yes | F | 3 | PuroR | 15 nt deletion (12 in intron) | del ex2-10 | 1:10 | Yes |
| GI-ME-N^Δ2-13^ C13 | Yes | F | 3 | PuroR | Del 28 nt exon 4 | del ex2-13 | 1:10 | Yes |
| GI-ME-N^Δ2-13^ C14 | Yes | F | 3 | PuroR | Del 28 nt exon 4 | del ex2-13 | 1:10 | Yes |
| GI-ME-N^Δ2-13^ C15 | Yes | F | 3 | PuroR | Del 28 nt exon 4 | del ex2-13 | 1:10 | Yes |
| GI-ME-N^Δ2-13^ C16 | Yes | F | 3 | PuroR | Del 28 nt exon 4 | del ex2-13 | 1:10 | Yes |
| NB139^Δ2-10^ C36 | Yes | M | 1 | del ex2-10 | - | - | 1:2 | Yes |
| NB139^Δ10-12^ C1 | Yes | M | 1 | del ex10-12 | - | - | 1:3 | Yes |
| NB139^Δ10-12^ C4 | Yes | M | 1 | del ex10-12 | - | - | 1:3 | Yes |
| SY5Y^-/-^ | No | F | 2 |  |  |  |  |  |
| SY5Y^Δ2-10^ | No | F | 2 |  |  |  |  |  |
| AMC753^-/-^ | No | M | 1 |  |  |  |  |  |

**Supplementary table 2. The number of significantly downregulated and upregulated differentially expressed genes for all 8 analyses with an p-adjusted value of 0.05.**

| Model | Number of downregulated genes | Number of upregulated genes |
| --- | --- | --- |
| SK-N-AS^-/-^ | 1982 | 1043 |
| GI-ME-N^-/-^ | 4000 | 4262 |
| GI-ME-N^Δ2-10^ | 3334 | 3568 |
| GI-ME-N^Δ2-13^ | 2748 | 3458 |
| NB139^Δ2-10^ | 3463 | 3205 |
| NB139^Δ10-12^ | 947 | 388 |
| PD^ΔATRX^ | 96 | 167 |
| iTHER^Δ2-10^ | 38 | 89 |

**Supplementary table 3. All media and their components used for culturing the cell lines/organoids and isogenic clones.**

| Cell line/Organoid | Main medium components | Supplements |
| --- | --- | --- |
| AMC772T2 | Dulbecco’s Modified Eagle’s Medium (DMEM) low glucose + GlutaMAX containing 1 g/L D-glucose and pyruvate (21885025, Gibco, Thermo Fisher) | 20% Ham’s F-12 Nutrient Mix + GlutaMAX (31765027, Gibco, Thermo Fisher), 1% Penicillin/Streptomycin (P/S, 15140122, Gibco, Thermo Fisher), B-27 without vitamin A (50X, 12587010, Gibco, Thermo Fisher), N2 supplement (100X) (17502048, Gibco, Thermo Fisher), 0.2 µg/ml Epithelial Growth Factor (hEGF), 0.4 µg/ml Fibroblast Growth Factor basic (hFGF), 2 µg/ml human Insulin-like Growth Factor 1 (hIGF1), and 0.1 µg/ml human Platelet Derived Growth Factor (PDGF) AA and BB. |
| CHLA90 and clones | Iscove’s Modified Dulbecco’s  Medium (IMDM) containing L-glutamine and 25 mM HEPES (12440053, Gibco, Thermo Fisher) | 10% Fetal Bovine Serum (FBS) (F0804, Sigma-Aldrich), 1% P/S, Insulin Transferrin Selenium (100X) (ITS, 41400045, Gibco, Thermo Fisher), and 2 mM L-glutamine (25030024, Gibco, Thermo Fisher). |
| GI-ME-N/GI-ME-N clones, SK-N-AS/SK-N-AS clones, SK-N-SH, LAN-6 and SJNB-12 | DMEM containing 4.5 g/L D-glucose, and L-glutamine (41965039, Gibco, Thermo Fisher) | 10% FBS, 1% P/S, 2 mM L-glutamine, and 1% MEM Non-Essential Amino Acids Solution (100X) (NEAA, 11140035, Gibco, Thermo Fisher). |
| NB139 motherline (TIC medium) | DMEM low glucose + GlutaMAX containing 1 g/L D-glucose and pyruvate (21885025, Gibco, Thermo Fisher) | 20% Ham’s F12 Nutrient Mix, 1% P/S, B-27 without vitamin A (50X), 0.2 µg/ml Epithelial Growth Factor (hEGF), and 0.4 µg/ml Fibroblast Growth Factor basic (hFGF). |
| NB139 clones (Organoid medium with 20% human plasma) | DMEM low glucose + GlutaMAX containing 1 g/L D-glucose and pyruvate | 20% Ham’s F-12 Nutrient Mix + GlutaMAX, 1% Penicillin/Streptomycin, B-27 without vitamin, N2 supplement (100X), 0.2 µg/ml Epithelial Growth Factor (hEGF), 0.4 µg/ml Fibroblast Growth Factor basic (hFGF), 2 µg/ml human Insulin-like Growth Factor 1 (hIGF1), and 0.1 µg/ml human Platelet Derived Growth Factor (PDGF) AA and BB. Additionally, we used 20% human plasma (add 5 ml MilliQ to one bottle of human plasma, P9523-5ML, Sigma), which solidifies in culture. |
| SKNMM | Roswell Park Memorial Institute (RPMI) 1640 medium containing L-glutamine (21875059, Gibco, Thermo Fisher) | 10% FBS and 1% P/S. |

**Supplementary table 4. Sequences of sgRNAs used to make isogenic knock-out and ATRX MEDs.** The guide efficiency is shown as predicted by the CRISPOR design tool.

| sgRNA name | sgRNA sequence 5' 🡪 3' | predicted guide efficiency |
| --- | --- | --- |
| ATRX_KO_sgRNA_1 | TCGTGACGATCCTGAAGACT | 81% |
| ATRX_KO_sgRNA_2 | CAGGATCGTCACGATCAAAG | 90% |
| ATRX_del2-10_sgRNA_1A | CACTACTATCGTATTACTTG | 88% |
| ATRX_del2-10_sgRNA_1B | TCGCCATTCTAACCGGCGTG | 98% |
| ATRX_del2-13_sgRNA_1A | TCGCCATTCTAACCGGCGTG | 98% |
| ATRX_del2-13_sgRNA_1B | TAGTGTTTATTAGAGTACGA | 88% |
| ATRX_del10-12_sgRNA_1A | TGGATTAGCCTACCCTGATG | 85% |
| ATRX_del10-12_sgRNA_1B | CAATACTATCAACTAAGCCG | 91% |

**Supplementary table 5. Primers used for cloning homology arm plasmids and PiggyBac doxycycline inducible *ATRX* wild-type plasmid.** Underlined sequences are homologous to the target sequence.

| primer name | primer sequence (5' --> 3') |
| --- | --- |
| ATRX_exon4_3'_homolgy_arm_1_forward | CTTCAGGATCGTCACGATCA |
| ATRX_exon4_3'_homolgy_arm_1_reverse | TCTGACAACTGGAACAATACCCA |
| ATRX_exon4_5'_homolgy_arm_1_forward | ATAGGGAGAGCGGCCGCGTCGAAGGCTACTGATACCCAG |
| ATRX_exon4_5'_homolgy_arm_1_reverse | GAAGATCTGGCGGCCGCGGATTTTTCTGAAGAGCTAGTTC |
| ATRX_exon4_3'_homolgy_arm_2_forward | AAGAGGTTGGTTGATGTTGAA |
| ATRX_exon4_3'_homolgy_arm_2_reverse | TCTGACAACTGGAACAATACCCA |
| ATRX_exon4_5'_homolgy_arm_2_forward | ATAGGGAGAGCGGCCGCGTCGAAGGCTACTGATACCCAG |
| ATRX_exon4_5'_homolgy_arm_2_reverse | GAAGATCTGGCGGCCGCTGATCGTGACGATCCTGAAGA |
| SalI_Kozak_ATRX_cDNA_FW | CGAGGTCGACGCCACCATGGCCGCTGAGCCCATG |
| ATRX_cDNA _RV2 | TGAACTAGTCTTCTTTGGAGAAAATCTG |
| XhoI_attL1_mCMV_FW | AGATCTCGAGCAAATAATGATTTTATTTTGACTGATAGTGACCTGTTCGTTGCAACAAATTGATAAGCAATGCTTTTTTATAATGCCAACTTTGTACAAAAAAGCAGGCTAAAGTGAAAGTCGAGCTC |
| SalI_mCMV_RV | TACCGTCGACTGATATCTGCAGAATTCCAC |
| BstEII_ATRX_FW | CACTGGGTCACCTCATGA |
| MluI_Att2L_RV | TTTACGCGTCAAATAATGATTTTATTTTGACTGATAGTGACCTGTTCGTTGCAACAAATTGATAAGCAATGCTTTCTTATAATGCCAACTTTGTACAAGAAAGCTGGGTTAAGATACATTGATGAGTTTGG |

**Supplementary table 6. Primers used for validation of patient-derived and isogenic models.** Primers with a matching sequence are coloured with a similar colour and were used for validating different aberrations.

| Primer name | Primer sequence (5' 🡪 3') | DNA or  RNA | Used on: | Confirms: |
| --- | --- | --- | --- | --- |
| check_puro_insert_In2_FW | GGAGGGGTTAGTATGCAAAAGG | DNA | GI-ME-N^-/-^; GI-ME-N^Δ2-10^; GI-ME-N^Δ2-13^ and SK-N-AS^-/-^ | presence PGK-eGFP-PuroR casette |
| check_puro_insert_PGK_RV | CTGCTAAAGCGCATGCTCCA |  |  |  |
| (WT/indel)_check_In3_FW | GGTTCTGGAAGTAACTCTGATATGA | DNA | All models | absence wildtype or presence indels in exon 4 |
| (WT/indel)_check_In4_RV | CATACTTTGTTACAATTGAAGGTTTC |  |  |  |
| SK-N-MM_K1367*_check_In11_FW | CGAGGCATTTTAAAGGCTGA | DNA | SK-N-MM | validate K1367* mutation in SK-N-MM |
| SK-N-MM_K1367*_check_In12_RV | CCACCACACCCAGCTATTTC |  |  |  |
| AMC772T2/CHLA-90_cDNA_UTR_FW | TGCATTTCTATCGTAACCGGG | RNA | AMC772T2;CHLA-90 | validate MED |
| AMC772T2/CHLA-90_cDNA_Ex14_RV | CTTCCAACTCTGCTTTCTTTGCAGAC |  |  |  |
| SK-N-MM_cDNA_Ex1_FW | ATGACCGCTGAGCCCATGAG | RNA | SK-N-MM | validate MED |
| SK-N-MM_cDNA_Ex14_RV | CTTCCAACTCTGCTTTCTTTGCAGAC |  |  |  |
| (WT/indel)_check_cDNA_Ex2_FW | GAAAGCAAGTTGAATACATTGGTGC | RNA | GI-ME-N^-/-^;SK-N-AS^-/-^ | absence wildtype or presence indels in exon 4 |
| (WT/indel)_check_cDNA_Ex6_RV | ATCATCTTTGTCTTCATTCAGCACTG |  |  |  |
| (WT/indel)_check_cDNA_Ex1_FW | ATGACCGCTGAGCCCATGAG | RNA | GI-ME-N^Δ2-10^ | absence wildtype or presence indels in exon 4 |
| (WT/indel)_check_cDNA_Ex6_RV | ATCATCTTTGTCTTCATTCAGCACTG |  |  |  |
| Ex2-10_MED_check_cDNA_Ex1_FW | ATGACCGCTGAGCCCATGAG | RNA | GI-ME-N^Δ2-10^; NB139^Δ2-10^ | validate exon 2-10 MED |
| Ex2-10_MED_check_cDNA_Ex12_RV | GCAAAAGCCTATGTCTGTATCTTG |  |  |  |
| Ex2-13_MED_check_cDNA_Ex1_FW | ATGACCGCTGAGCCCATGAG | RNA | GI-ME-N^Δ2-13^ | validate exon 2-13 MED |
| Ex2-13_MED_check_cDNA_Ex15_RV | CCAGGAGACTTGGAATCATCATTTTC |  |  |  |
| Ex10-12_MED_check_cDNA_Ex9_FW | GGTGAGGGAAGCAGCGATGAAC | RNA | NB139^Δ10-12^ | validate exon 10-12 MED |
| Ex10-12_MED_check_cDNA_Ex15_RV | CCAGGAGACTTGGAATCATCATTTTC |  |  |  |

**Supplementary table 7. Antibodies used for western blotting and for immune fluorescence.**

| Target | Antibody type | Western blot dilution | IF dilution | Supplier |
| --- | --- | --- | --- | --- |
| ATRX | Rabbit pAb, IgG | 1:500 | 1:50 | Abcam (ab97508) |
| COX-IV | Rabbit mAb, IgG | 1:2000 |  | Abcam (ab202554) |
| HA-tag | Rabbit mAb, IgG | 1:250 |  | Cell Signaling (C29F4) |
| Histone H3.3 | Rabbit pAb, IgG | 1:500 |  | Merck Millipore (09-838) |
| mH2A1 | Rabbit pAb, IgG | 1:1000 |  | Abcam (ab37264) |
| α-tubulin | Mouse mAb, IgG | 1:10,000 |  | Cell Signaling (DM1A) |
| γH2A.X | Rabbit pAb, IgG | 1:500 |  | Abcam (ab2893) |
| Mouse IgG | Sheep IgG-HRP | 1:5000 |  | Amersham ECL GE Healthcare (NXA931V) |
| Rabbit IgG | Donkey IgG-HRP | 1:5000 |  | Amersham ECL GE Healthcare (NA9340V) |
| Mouse IgG | Goat IgG IRDye® 680RD | 1:5000 |  | LI-COR (926-68070) |
| MYCN | Mouse mAb, IgG | 1:500 |  | Santa Cruz Biotechnology (sc-53993) |
| GAPDH | Rabbit mAb, IgG | 1:1000 |  | Cell Signaling (2118S) |
| cMYC | Rabbit mAb, IgG | 1:1000 |  | Cell Signaling (5605S) |
| EZH2 | Rabbit mAb, IgG | 1:1000 |  | Cell Signaling (5246S) |
| HP1α | Mouse mAB, IgG |  | 1:500 | Merck Millipore (05-689) |
| PML | Mouse mAb, IgG |  | 1:50 | Abcam (ab96051) |
| Mouse IgG | Donkey IgG Alexa fluor 647 |  | 1:50 | ThermoFisher (A-31571) |
| Rabbit IgG | Donkey IgG Alexa fluor 647 |  | 1:50 | ThermoFisher (A-A31573) |
| Mouse IgG | Donkey IgG Alexa fluor 488 |  | 1:50 | ThermoFisher (A-21202) |

**Supplementary table 8. Primers used for 47S rRNA qPCRs.**

| Primer name | Primer sequence (5' 🡪 3') |
| --- | --- |
| qPCR_rRNA_ITS2_FW1 | TCCCGAGCTTCCGCGTCG |
| qPCR_rRNA_ITS2_RV1 | GACCGGAGGGAGGGGCAC |
| qPCR_UBE3B_FW_1 | ACCTCGAGAGCATGGTTCATC |
| qPCR_UBE3B_RV_1 | CAGTCGACTCCGACAGAGAAA |
| qPCR_ATP5PO_FW_1 | TCCTGAAGGAACCCAAAGTGG |
| qPCR_ATP5PO_RV_1 | TGATCAGATTGGTAGTGAGGGG |
